# Stromal estrogen signaling regulates fallopian tube homeostasis and cancer initiation via inflammatory pathways

**DOI:** 10.64898/2026.01.21.700864

**Authors:** Xueqing Chen, Sen Han, Dongyi Zhao, Guyu Qin, Zhe Li

## Abstract

Lifetime estrogen exposure is a major risk factor for ovarian cancer, which can originate from fallopian tube epithelial (FTE) cells. Here we report that estrogen receptor α (ERα) signaling in fallopian tube (FT) stromal cells plays a critical developmental role in maintaining epithelial homeostasis by promoting FTE proliferation and ciliated differentiation. Stromal ERα regulates expression of inflammatory cytokines, growth factors, and extracellular matrix components, creating a differentiation-supportive, tumor-suppressive niche that coordinates epithelial regeneration during hormonal cycles. Excessive or prolonged estrogen exposure, however, shifts this niche toward a tumor-promoting state by inducing inflammation and activating stemness-associated pathways, including JAK/STAT, in FTE cells. This effect is exacerbated in genetically altered FTE cells lacking key tumor suppressors, which resist differentiation while remaining responsive to stromal proliferation signals. These findings reveal how stromal ERα signaling integrates hormonal cues, inflammation, aging, and genetic susceptibility to influence early events in FT carcinogenesis.

**In Brief:** Estrogen receptor α signaling in fallopian tube stromal cells maintains epithelial homeostasis by promoting proliferation and ciliated differentiation. Excessive estrogen shifts this niche toward inflammation and stemness, cooperating with genetic susceptibility to drive early fallopian tube carcinogenesis.

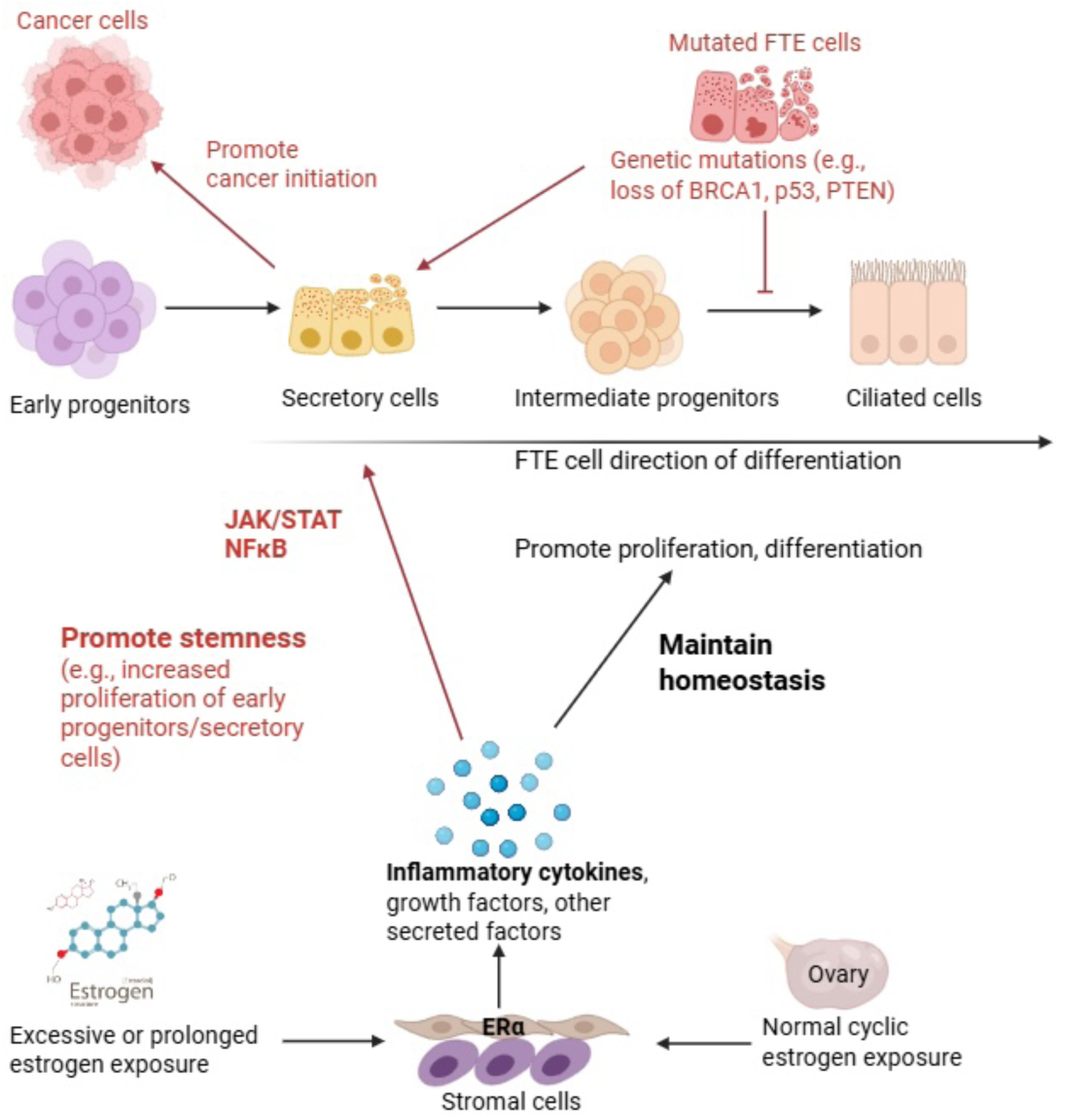

**Highlights:**

- Stromal ERα signaling maintains fallopian tube epithelial (FTE) cell homeostasis
- Stromal ERα promotes proliferation and ciliated differentiation of FTE cells
- Excess estrogen shifts this stromal niche toward tumor promotion via inflammation
- Differentiation-resistant mutant FTE cells respond to stromal proliferation signals

## Introduction

Epithelial ovarian cancer (EOC) accounts for more than 95% of ovarian malignancies and is the most lethal gynecological cancer among women worldwide ^1,2^. In addition to genetic risk factors such as inherited pathogenic mutations in *BRCA1/2* or other high-penetrance genes, several non-genetic factors can also increase the risk of EOC. Among these environmental and lifestyle risk factors, long-term exposure to elevated estrogen levels is well documented. Strong epidemiological evidence supports that the etiology, pathogenesis, and progression of EOC are highly dependent on estrogen signaling activities ^3,4^. Prolonged use of unopposed menopausal hormone therapy (i.e., estrogen without progesterone) is associated with an increased risk of EOC ^5–7^. Moreover, greater lifetime cumulative exposure to estrogen, resulting from factors such as early menarche, late menopause, and nulliparity, may also elevate EOC risk ^8–10^. Despite these associations, the mechanisms by which estrogen contributes to EOC development remain poorly understood. Elucidating the molecular and cellular effects of estrogen may provide critical insights for EOC risk reduction and prevention strategies.

Although EOCs were initially thought to originate from ovarian surface epithelial cells, accumulating evidence now indicates a fallopian tube (FT) origin for most cases. Pathological analyses of human precancerous lesions, such as serous tubal intraepithelial carcinoma (STIC) identified in the fimbrial region of the FT ^11,12^, together with genomic studies of patient samples and genetically engineered mouse models, strongly support the conclusion that most EOCs, particularly high-grade serous ovarian cancer (HGSOC), arise from fallopian tube epithelial (FTE) cells ^13–17^. The FT epithelium consists of PAX8⁺OVGP1⁺ secretory cells and acetylated tubulin (AcTUB)⁺ ciliated cells. Lineage-tracing studies in mouse oviducts, as well as single-cell analyses of human FT tissues, demonstrate that secretory cells possess stem and progenitor cell properties and can differentiate into ciliated cells, which represent a more differentiated FTE lineage ^18–20^. Accordingly, most HGSOCs are thought to originate from FTE secretory cells, particularly those located in the fimbrial (distal) region of the FT adjacent to the ovary ^11,12^. Despite these advances, the molecular and cellular mechanisms driving HGSOC initiation from FTE secretory cells remain poorly understood.

The specification and maintenance of epithelial cells and their stem/progenitor cells are dominantly controlled by their supporting cells, or niche, which secrete factors that regulate epithelial stem cell self-renewal and differentiation through paracrine signaling ^21–23^. Disruption of this regulatory control can promote malignant transformation ^23^. In particular, chronic inflammation within the niche may contribute to the establishment of a cancer-promoting microenvironment ^24^. In the FT, stromal cells are integral components of the local microenvironment and play essential roles in maintaining FTE homeostasis, supporting reproductive function, and influencing disease susceptibility, including ovarian cancer.

In a previous study ^25^, we identified a stromal subpopulation in the mouse oviduct (i.e., the mouse equivalent of the human FT) enriched for *Pdgfra*^+^ and *Esr1*^+^ cells, where *Esr1* encodes estrogen receptor α (ERα). These stromal cells express multiple secreted factors associated with developmental and growth pathways that can regulate the proliferation and differentiation of adjacent FTE cells via paracrine signaling, suggesting their role as a functional niche for FTE homeostasis. To investigate the functional significance of this estrogen-responsive stromal niche, we performed a conditional knockout study of *Esr1* in *Pdgfra*^+^ stromal cells *in vivo* coupled with *in vitro* experiments using an FTE cell/FT stromal cell co-culture organoid system. Here we report that estrogen/estrogen receptor (ER) signaling in stromal cells is a key regulator of FT homeostasis, in part through activation of inflammation pathways that drive FTE cell proliferation and differentiation. Importantly, excessive estrogen exposure may promote EOC initiation by disrupting FTE homeostasis through dysregulation of FTE cells via this stromal niche.

## Results

### Stromal estrogen/ER signaling regulates FT homeostasis

Since *Esr1^+^* FT stromal cells residue within the CD140a^+^ subpopulation (CD140a is encoded by *Pdgfra*) ^25^, we used the *Pdgfra-Cre* mouse line^26^ to disrupt *Esr1* conditional knockout alleles (*Esr1^fl/fl^*) in stromal cells (Figure 1A). In a subset of mice, the conditional Cre-reporter *Rosa26-LSL-YFP* (*R26Y*)^27^ was included to genetically mark stromal cells undergoing *Pdgfra-Cre*-mediated recombination. In the resulting *Pdgfra-Cre;Esr1^fl/^*^fl^*;R26Y* (*PEY*) female mice, the oviducts (FTs) were notably smaller than those of *Pdgfra-Cre;R26Y* (*PcY*) control female mice (Figure 1B). This reduction in FT size was evident in *PEY* females as early as ∼4 weeks of age (Figure S1A). Since *Pdgfra-Cre* is also expressed in stromal cells of other organs, we assessed whether the observed FT phenotype could be attributed to systemic alterations in ovarian hormone levels, such as estrogen. Serum estradiol (E2) levels were measured across the estrous cycle in *PEY* and *PcY* females, and no significant differences were detected (Figure S1B), thus largely ruled out this possibility.

**Figure 1.**
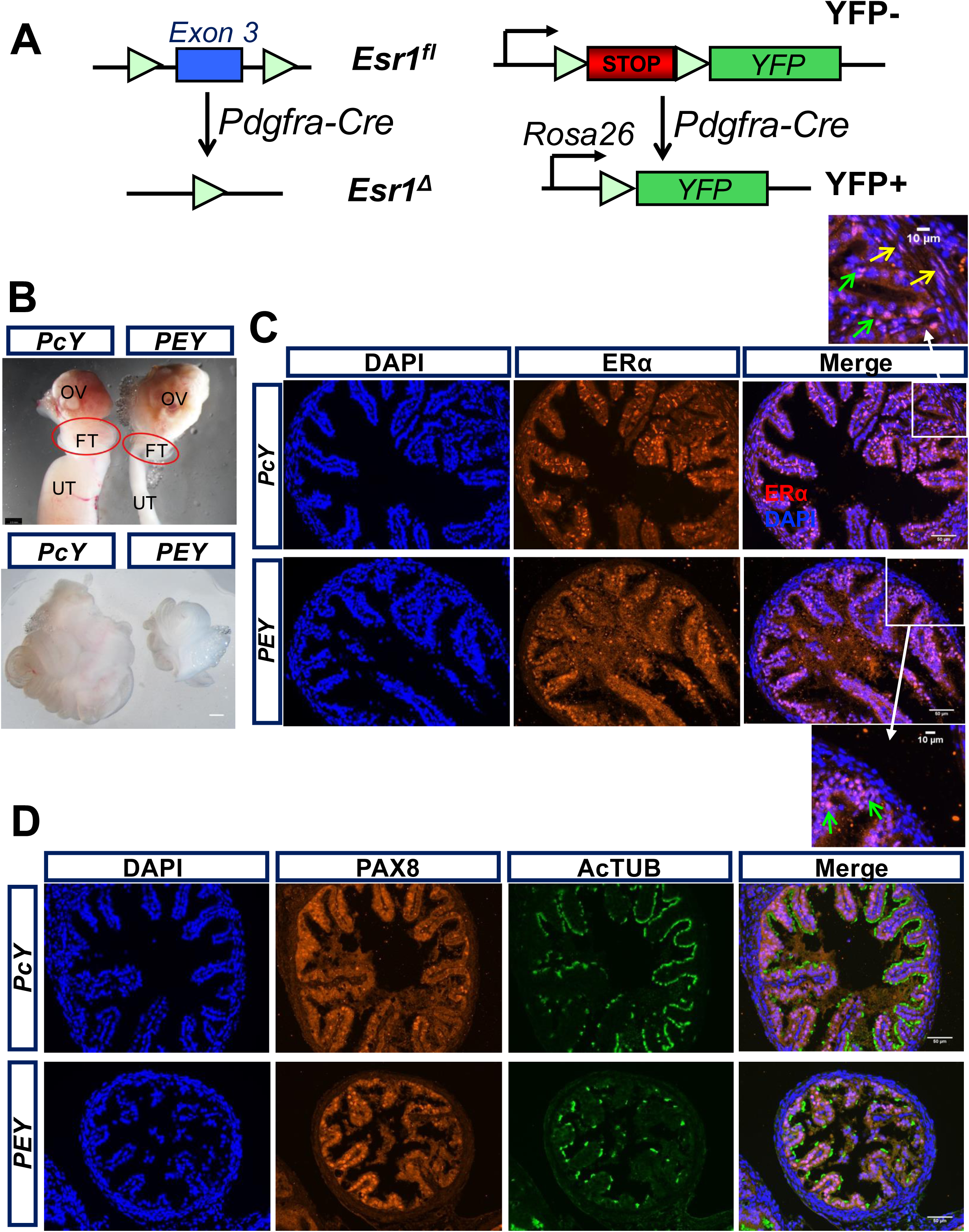
*Pdgfra-Cre*-induced *Esr1*-loss leads to impaired FT homeostasis. (A) Schematic diagram of conditional knockout (CKO) of *Esr1* driven by *Pdgfra-Cre*; cells with *Esr1*-loss are genetically marked by YFP. (B) Dissection of ovary (OV), fallopian tube (FT) and uterus (UT) from adult (∼8 weeks) female mice with the genotype of *Pdgfra-Cre;R26Y* control (*PcY*) or *Pdgfra-Cre;Esr1^fl/fl^;R26Y* CKO (*PEY*). Scale bars=2.5 mm. (C) Immunofluorescence (IF) staining of FT sections from mice with the indicated genotypes for Estrogen Receptor alpha (ERα); in the insets, yellow arrows indicate FT stromal cells positive for ERα, green arrows indicate FTE cells positive for ERα. Scale bars= 50μm (10μm in the insets). (D) Co-IF staining of FT sections from mice with the indicated genotypes for Paired Box 8 (PAX8) and Acetyl-alpha Tubulin (AcTUB). Scale bars= 50μm.

In FT sections from *PEY* and *PcY* mice, immunofluorescence (IF) staining for ERα revealed that while ERα expression was preserved in FTE cells in both genotypes, its expression was specifically lost in FT stromal cells of *PEY* mice (Figure 1C, showing fimbrial region). The reduced FT size observed in *PEY* mice suggests that loss of *Esr1* in stromal cells affects not only the stromal compartment but also the epithelial layer. To assess potential effects on FTE cells, we performed co-IF staining of FT sections for FTE lineage markers, including the secretory cell maker PAX8 and the ciliated cell marker AcTUB. Notably, FTE cells in *PEY* mice exhibited reduced AcTUB staining compared with *PcY* controls (Figure 1D), indicating that stromal *Esr1*-loss may impair ciliated cell differentiation.

To further characterize FTE cells, we employed an organoid culture platform we established previously for murine FTE cells ^28^. Organoid assays revealed that fewer organoids formed per FT from *PEY* mice, consistent with their reduced FT size, and that the resulting organoids were also smaller than those derived from *PcY* controls (Figure S1C-D). When seeded at equal cell numbers, *PEY*-derived cultures generated comparable numbers of organoids; however, these organoids remained smaller (Figure S1E-F). These findings indicate alterations in FTE stem/progenitor cell properties, potentially including impaired differentiation toward the ciliated lineage, resulting in the formation of smaller and less mature organoids. Despite the reduced FT size, *PEY* females remained fertile and were able to produce at least one successful litter, although uterine abnormalities may constrain subsequent pregnancies. Collectively, these data demonstrate that stromal estrogen/ER signaling is critical for FT homeostasis and that loss of *Esr1* in FT stromal cells exerts profound effects on both stromal and epithelial compartments.

### Stromal *Esr1* expression is required for stromal niche-mediated regulation of FTE cells

We previously reported that FT stromal cells function as a niche for FTE cells by producing multiple secreted factors ^25^. To further elucidate how FT stromal cells regulate FTE cells, particularly secretory cells, we sorted Lin^-^CD24^+^ FTE cells from freshly dissected FTs of wild-type (WT) adult female mice and cultured them in our previously defined BET organoid culture medium, composed of <u>B</u>27, <u>E</u>GF, and <u>T</u>GFBR1 inhibitor ^28^. Under this condition, FTE cells are maintained long-term as secretory cells upon serial passaging ^25^. We then co-cultured established FTE organoid cells (i.e., secretory cells) with freshly sorted Lin^-^CD24^-^ FT stromal cells from WT mice (Figure S2A). 7 days after co-culture, EpCAM^+^ epithelial cells were isolated from the resulting organoids (FTE+SC) and compared with EpCAM^+^ cells from matched organoid cultures without stromal cells (FTE-only) by RNA sequencing (RNA-seq). Gene-set enrichment analysis (GSEA)^29^ of the RNA-seq data revealed that the top-enriched gene sets in the Hallmark collection of the Molecular Signatures Database (MSigDB) were associated with cell cycle regulation (Figure 2A), suggesting stromal cell-induced proliferation of FTE cells.

**Figure 2.**
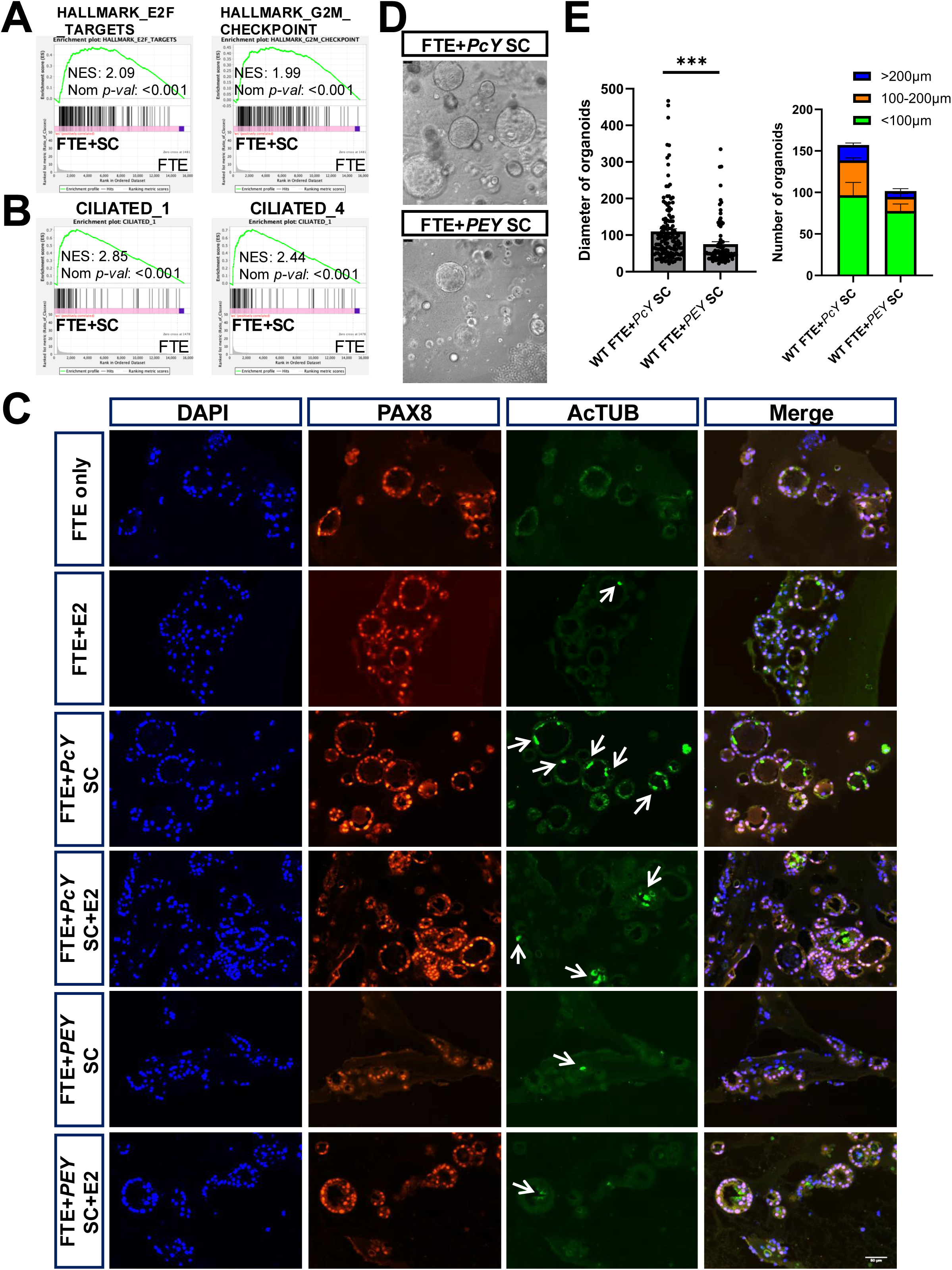
Stromal *Esr1* is required for the regulation of FTE growth by stromal cells. (A-B) GSEA showing enrichment of the indicated Hallmark [from the Molecular Signatures Database (MSigDB)] gene sets (A) or gene sets representing different human FTE subsets^19^ (B) in FTE organoid cells sorted from the FTE cell/stromal cell co-culture (FTE+SC) in relation to those from the FTE cell-only culture (FTE). NES: normalized enrichment score; Nom p-val: nominal p value. (C) Co-IF staining of organoid sections for PAX8 and AcTUB. Organoids were from the indicated culture condition (SC: stromal cells from either *PcY* control or *PEY* CKO mice; E2: 1nM). Arrows indicate organoid cells with apical AcTUB staining. Scale bars=50μm. (D) Representative pictures from established wild-type (WT) FTE organoids co-cultured with FT stromal cells sorted from *PcY* or *PEY* mice. Scale bars=50μm. (E) Quantification of sizes and numbers (at different size ranges) of organoids formed from different organoid cultures as in (D). ***: p≤0.001, Student’s t-test. Error bar represents ±SEM.

In a recent single-cell transcriptomic study of human FTE cells (*Dinh et al* ^19^), pseudotime analysis identified early progenitor populations that differentiate into secretory cells and subsequently into ciliated cells through intermediate progenitors. Using these human FTE lineage signatures as gene sets, we found that multiple ciliated cell-related gene sets were significantly enriched in FTE cells co-cultured with stromal cells (FTE+SC, Figure 2B). Consistently, gene ontology (GO) biological process (BP) terms related to ciliated cell function were among the top enriched gene sets in FTE cells from the FTE+SC cultures (Figure S2B, highlighted). These data suggest that FT stromal cells promote both proliferation and ciliated differentiation of FTE cells derived from secretory populations. Co-IF staining of organoid sections further confirmed the increased presence of differentiated AcTUB^+^ ciliated cells and proliferative Ki67^+^ cells in FTE+SC organoids compared with FTE-only organoids (Figures 2C and S2C, FTE+*PcY* SC). In addition, quantitative RT-PCR (qRT-PCR) analysis showed upregulation of *Foxj1*, a key transcription factor required for ciliated cell differentiation ^30^, in organoids from the FTE+*PcY* SC co-cultures (Figure S2D).

Next, to determine whether *Esr1* expression in FT stromal cells is required for this stromal regulation of FTE cells, we isolated Lin^-^CD24^+^ stromal cells from FTs of *PEY* mice or *PcY* control mice and co-cultured them with established WT FTE organoid cells. Co-culture with *Esr1*-deficient stromal cells from *PEY* mice resulted in the formation of significantly fewer and smaller organoids compared with co-cultures containing *PcY* stromal cells (Figure 2D-E). Co-IF staining of their corresponding organoid sections demonstrated that the *PcY* stromal cell-induced increase of the AcTUB^+^ ciliated cells and Ki67^+^ proliferative cells was largely abolished when *PEY* stromal cells were used (Figures 2C and S2C, FTE+*PEY* SC). Consistently, *Foxj1* expression was reduced in organoids from FTE+*PEY* SC co-cultures relative to those from FTE+*PcY* SC co-cultures (Figure S2D). Finally, co-culture of stromal cells isolated from *PEY* or *PcY* mice with freshly sorted Lin^-^CD24^+^ FTE cells from WT adult female mice yielded similar results, with a reduction in organoid formation in the presence of *PEY* stromal cells (Figure S2E-F). Collectively, these findings demonstrate that FT stromal cells support FTE proliferation and ciliated differentiation and that stromal *Esr1* expression is essential for this critical stromal-epithelial regulatory interaction.

### Stromal *Esr1* is linked to inflammatory pathways and secreted factors

To investigate how estrogen/ER signaling in FT stromal cells regulates FTE cells, we sorted YFP-labeled *Esr1*-null or *Esr1*-WT stromal cells from the FTs of *PEY* or *PcY* mice, respectively, and subjected them to RNA-seq. Among the most significantly downregulated genes in *Esr1*-null stromal cells were numerous genes associated with the extracellular matrix (ECM) (e.g., *Postn*, *Col5a3*) and inflammation/immune response (e.g., *Cxcl2*, *Il4ra*) (Figure S3A). Consistent with these findings, GSEA revealed that inflammation/immune-related signatures were among the most significantly downregulated Hallmark gene sets in *Esr1*-null stromal cells (Figure 3A, highlighted). This downregulation was largely attributable to reduced expression of many cytokine and cytokine receptor genes (Figure 3B). In addition, gene sets related to secreted factors and ECM organization were significantly downregulated in *Esr1*-null stromal cells (Figure 3C). Many downregulated secreted factors included not only inflammation-associated cytokine genes but also growth factor genes (Figure 3C). Among hormone receptor genes, deletion of *Esr1* resulted in reduced expression of *Pgr*, which encodes the progesterone receptor and is a well-established ERα target gene ^31^ (Figure S3B). Differential expression of selected genes identified by RNA-seq was validated by qRT-PCR in sorted YFP^+^ *Esr1*-null versus *Esr1*-WT FT stromal cells (Figure 3D). Notably, we previously reported that *Esr1^+^* FT stromal cells express *Igf1* and that its protein product, IGF1, promotes FTE proliferation and differentiation ^25^. However, our RNA-seq and validation analyses showed that *Igf1* expression was not significantly altered in FT stromal cells following *Esr1* deletion, suggesting that *Igf1* is not a direct ERα target gene in this cellular context (Figures 3D and S3B).

**Figure 3.**
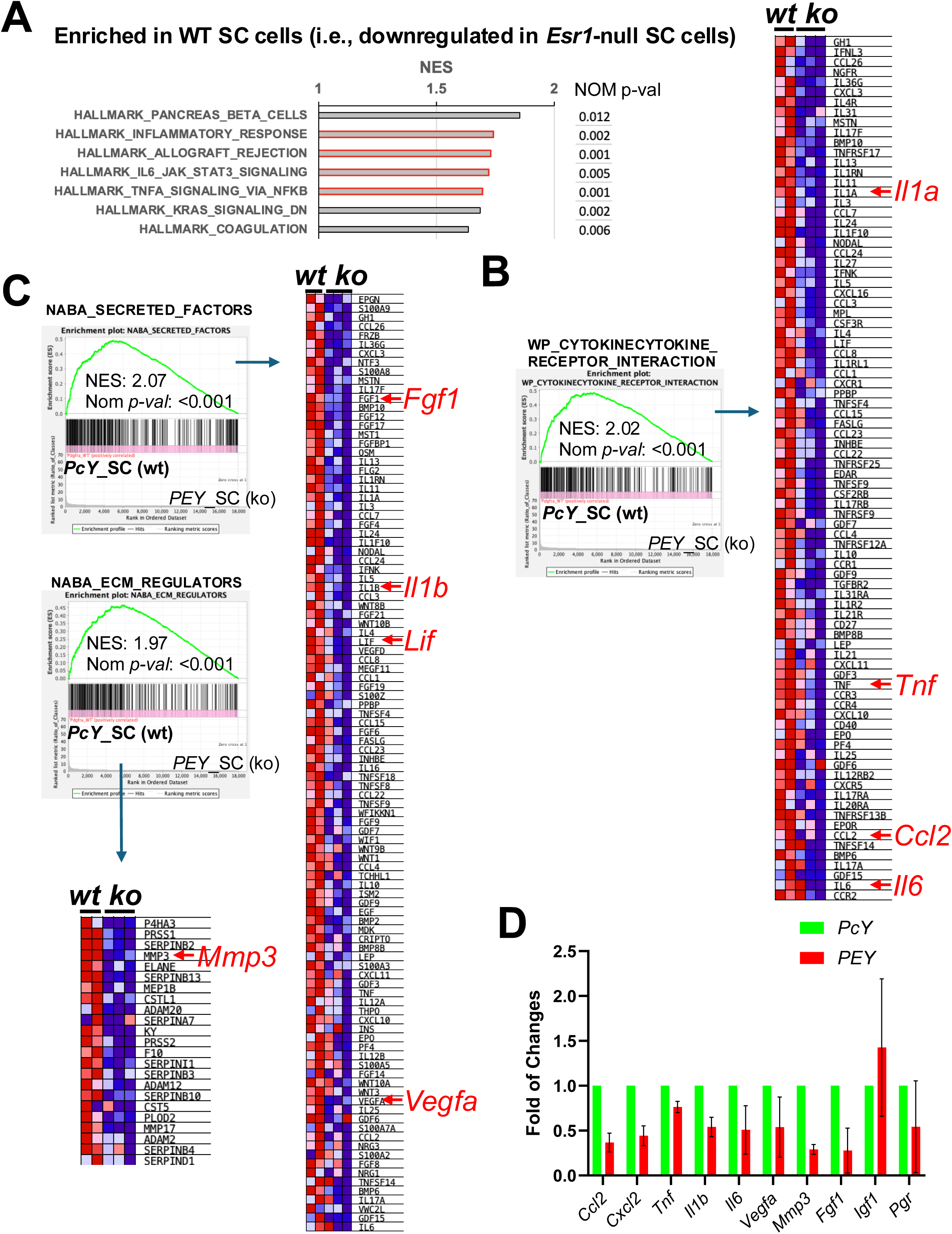
Stromal *Esr1* is linked with inflammatory pathways and secreted factors. (A) GSEA data showing top enriched gene sets in the Hallmark collection from the MSigDB in YFP^+^ FT stromal cells sorted from *PcY* control mice in relation to those from *PEY* (*Esr1*-CKO) mice. Gene sets related to inflammation/immune pathways are highlighted. (B-C) GSEA plots and heatmaps showing enrichment of representative cytokine/cytokine receptor-related (B) and secreted factor/ECM regulator-related (C) gene sets in *PcY* YFP^+^ stromal cells (*wt*) in relation to *PEY* YFP^+^ stromal cells (*ko*). In the heatmaps, red to blue represent highest to lowest expression, top significantly enriched genes are shown and representative genes related to inflammation cytokines, growth factors, and ECM are indicated. (D) Quantitative Reverse Transcription PCR (qRT-PCR) analysis of genes of interest in *PcY* or *PEY* YFP^+^ FT stromal cells. Error bar represents ±SEM.

Of note, *Esr1* is also expressed in FTE cells (e.g., Figure 1C) and epithelial-specific deletion of *Esr1* in the oviduct using *Wnt7a*-Cre has been shown to impair fertilization and embryo survival as a result of elevated secretion of innate immune mediators, including proteases ^32^. This phenotype and underlying molecular mechanism are distinct from those observed following stromal-specific *Esr1* deletion described here. Consistent with this distinction, retrospective analysis of gene expression changes related to inflammatory cytokines, ECM regulators/factors, secreted growth factors, and hormone receptors in *Esr1*-null versus WT oviducts from the epithelial-deletion study did not recapitulate the expression patterns observed in *PEY* versus *PcY* mice (Figure S3C, compared to Figure S3B). Together, these results indicate that estrogen signaling through ERα in FT stromal cells supports epithelial-stromal homeostasis by promoting the expression of proinflammatory cytokines, ECM components, and growth factors.

### Inflammatory cytokines and secreted growth factors regulate FTE growth

To directly test whether proinflammatory cytokines predicted to be produced by FT stromal cells can regulate FTE cells, we selected several cytokine genes that were significantly downregulated in *Esr1*-null stromal cells (e.g., *Tnf*, *Il1a*, *Il1b*, Figures 3B-C and S3B). Supplementation of the minimal BET organoid culture medium^28^ with TNFα, IL-1α, or IL-1β significantly increased both the number and size of WT FTE organoids (Figure 4A-B). IF staining of organoid sections for PAX8 and AcTUB revealed a marked increase in AcTUB^+^ organoid cells upon TNFα treatment, whereas IL-1α or IL-1β treatment resulted in only a modest increase in AcTUB staining compared with those in the BET medium alone (Figure S4). These findings suggest that inflammatory cytokines are capable of enhancing FTE growth, at least in part, by promoting ciliated differentiation, albeit to varying extents.

**Figure 4.**
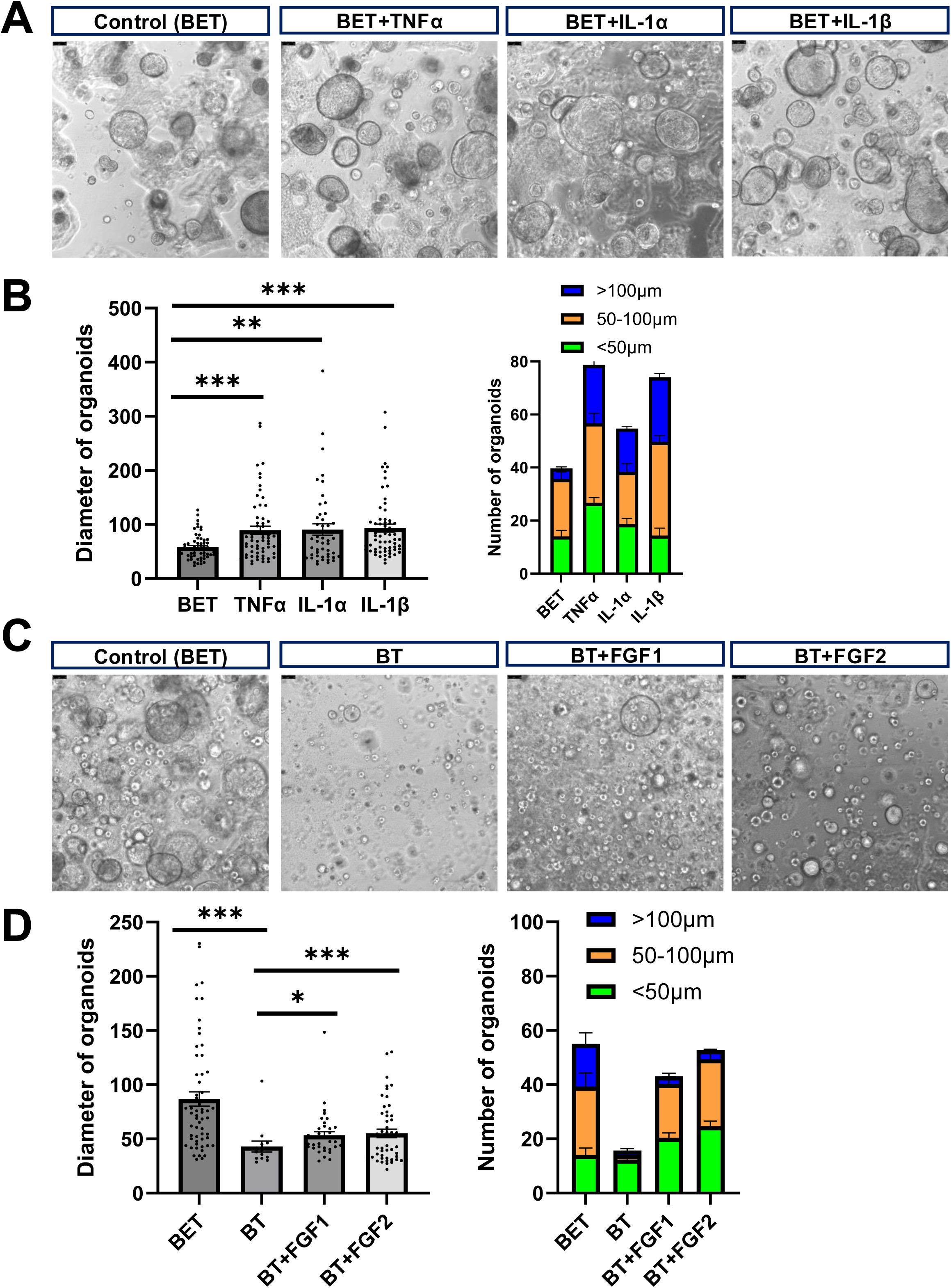
Stromal *Esr1*-linked inflammatory cytokines and secreted growth factors regulate FTE growth. (A) Representative pictures from organoid cultures for WT FTE cells growing in the BET medium supplemented with the indicated cytokines (10ng/μl). Scale bars=50μm. (B) Quantification of sizes and numbers (at different size ranges) of organoids formed from different cultures as in (A). **: p≤0.01, ***: p≤0.001, Student’s t-test. Error bar represents ±SEM. (C) Representative pictures from organoid cultures for WT FTE cells growing in the BT medium supplemented with the indicated growth factors (10ng/μl) or in the BT or BET medium (as controls). Scale bars=50μm. (D) Quantification of sizes and numbers (at different size ranges) of organoids formed from different cultures as in (C). *: p≤0.05, ***: p≤0.001, ANOVA. Error bar represents ±SEM.

We next examined the effects of stromal-derived growth factors, focusing on FGF1 and the related family member FGF2. Since inclusion of EGF in the BET medium may activate growth-promoting pathways that overlap with FGF/FGFR signaling, we removed EGF from the BET medium to generate the BT medium. Supplementation of BT medium with either FGF1 or FGF2 significantly increased both the number and size of WT FTE organoids, although the effects were less pronounced than those observed with EGF-containing BET medium (Figure 4C-D). Collectively, these results suggest that inflammatory cytokines and secreted growth factors regulated by stromal *Esr1* are capable of promoting FTE cell growth through paracrine signaling mechanisms.

### Aging-associated stromal CCL2 regulates FTE growth

Recent single-cell studies of the murine female reproductive tract (FRT) have shown that the FT (oviduct) undergoes continuous remodeling across the estrous cycle, resulting in chronic inflammation with aging ^33^. During the estrous cycle, estrogen levels peak prior to ovulation during proestrus (P), followed by increased progesterone levels during metestrus (M) and diestrus (D) ^34^. At ovulation, corresponding to estrus (E), fimbrial cells are exposed to follicular fluid containing locally high concentrations of estrogen ^35,36^. Analysis of published single-cell transcriptomic data^33^ for FT stromal cells across different phases of the estrous cycle revealed that in FT stromal cells, expression of estrogen signaling-related genes peaks at ovulation and coincides with maximal expression of inflammation-, cytokine-, and ECM-associated genes (Figure S5A). Notably, these same pathway genes are downregulated in *Esr1*-deficient FT stromal cells in *PEY* mice (Figures 3 and S3), thus further supporting a role for stromal estrogen/ER signaling in ECM remodeling and proinflammatory cytokine production.

The same single-cell study also demonstrated that FT aging is associated with increased immune cell infiltration, potentially driven by repeated ECM remodeling during the estrous cycle and incomplete resolution of estrogen signaling-induced inflammation ^33^. Consistent with this, aging markedly enhanced inflammatory signaling in FT stromal fibroblasts, and reanalysis of the dataset showed that several cytokine genes, including *Ccl2*, *Ccl7*, *Cxcl1*, and *Cxcl2*, were among the most significantly upregulated genes in FT stromal cells from aged (18-month-old) mice compared with young mice (Figure S5B). GSEA further confirmed immune-related gene sets as the most significantly enriched pathways in aged FT stromal fibroblasts (Figure S5C, highlighted).

Our RNA-seq analysis of *Esr1*-null versus *Esr1*-WT FT stromal cells revealed downregulation of these aging-associated cytokine genes in *Esr1*-deficient stromal cells (Figures 3D and S3B). We focused on one of them, *Ccl2*, for functional validation. In organoid culture, supplementation of BET medium with recombinant CCL2 significantly enhanced FTE organoid growth, resulting in increased organoid number and size (Figure 5A-C). Conversely, co-culture of WT FTE organoids with *Ccl2^-/-^* FT stromal cells (derived from *Ccl2* knockout mice^37^) led to a reduction in stromal support for FTE organoid growth compared with co-culture with WT stromal cells (Figure 5A-C). Co-IF staining of organoid sections demonstrated that CCL2 treatment increased the numbers of AcTUB^+^ ciliated cells and Ki67^+^ proliferative cells relative to those in BET medium alone, whereas co-culture with *Ccl2^-/-^* FT stromal cells resulted in reduced numbers of both AcTUB^+^ and Ki67^+^ cells compared with co-culture with WT stromal cells (Figures 5D and S5D). Together, these findings indicate that aging-associated, stromal *Esr1*-regulated cytokines, exemplified by CCL2, promote FTE cell proliferation and differentiation, thus linking estrogen/ER signaling-related stromal inflammation during aging to estrogen-regulated epithelial homeostasis in the FT.

**Figure 5.**
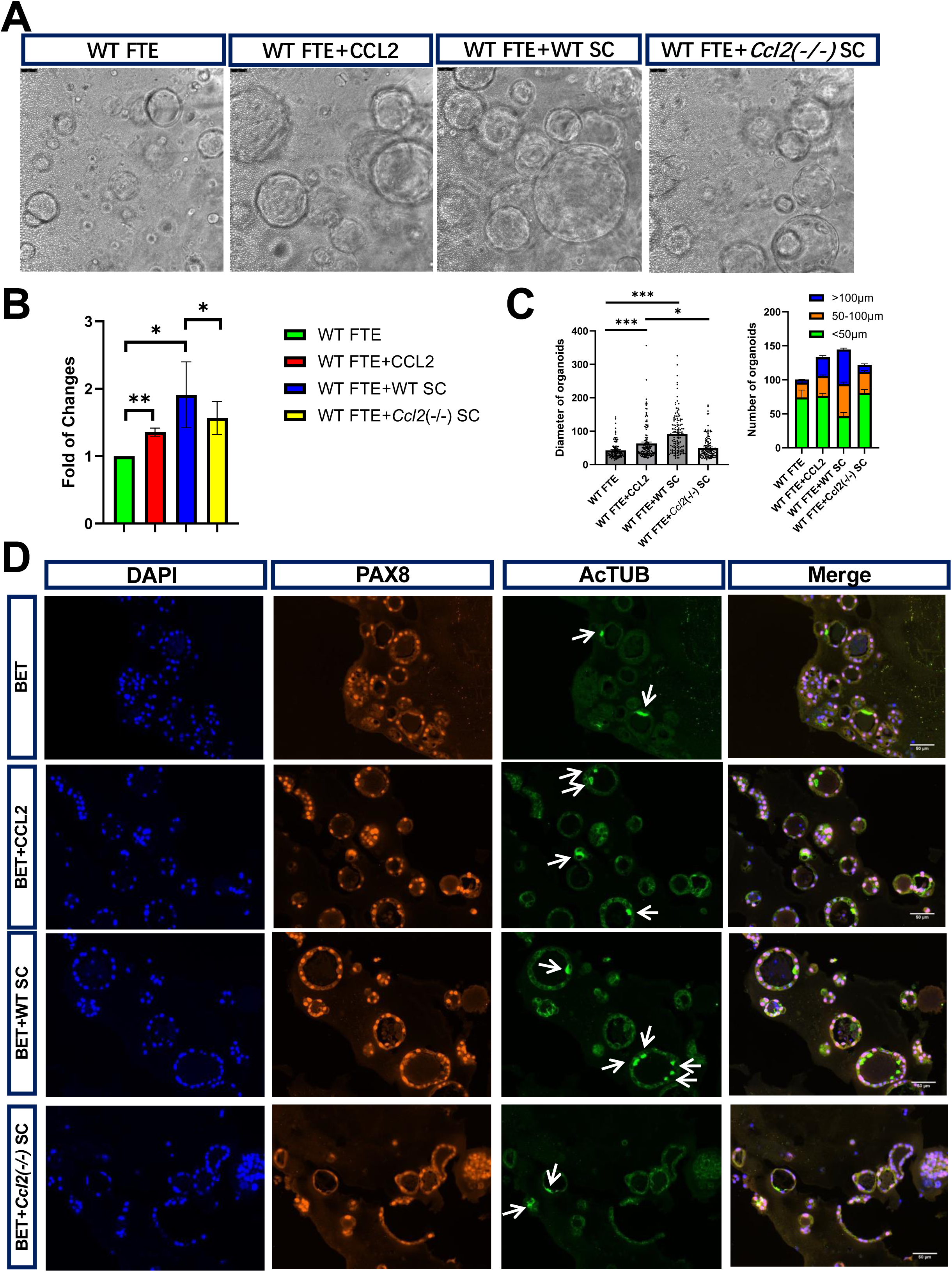
Aging-related stromal CCL2 regulates FTE growth. (A) Representative pictures from organoid cultures for WT FTE cells growing in the BET medium supplemented CCL2 (10ng/μl) or co-cultured with FT stromal cells MACS-sorted from WT or *Ccl2^-/-^* mice. Scale bars=50μm. (B) ATP assay of organoids in (A). Data was normalized to the values from WT FTE organoids (=1). *: p≤0.05, **: p≤0.01, Student’s t-test. Error bar represents ±SEM. (C) Quantification of sizes and numbers (at different size ranges) of organoids formed from different cultures as in (A). *: p≤0.05, ***: p≤0.001, Student’s t-test. Error bar represents ±SEM. (D) Co-IF staining of organoids sections for PAX8 and AcTUB. Organoids were from the indicated culture condition. Arrows indicate organoid cells with apical AcTUB staining. Scale bars= 50μm.

### Estrogen promotes inflammation and growth of FTE cells via *Esr1^+^* stromal cells

To directly examine how estrogen exposure influences FTE cells, we used the organoid culture and co-culture systems described above (Figure 2). Treatment of WT FTE organoids cultured in BET medium with increasing concentrations of estradiol (E2) did not result in significant changes in organoid size or morphology (Figure S6A). In contrast, while co-culture of WT FTE cells with WT FT stromal cells (FTE+SC) markedly enhanced organoid growth, addition of E2 (1nM) to this co-culture further increased both organoid size and number (FTE+SC+E2, Figure 6A-B). GSEA analysis of RNA-seq data from sorted FTE organoid cells cultured under these different conditions revealed that inclusion of FT stromal cells robustly induced upregulation of several human FTE lineage signatures defined by *Dinh et al* ^19^, with strongest enrichment of the ciliated cell signatures Ciliated_1 and Ciliated_4, followed by Unclassified_3 and Unclassified_1 (Figure 6C; FTE+SC, light blue line). Notably, addition of E2 to the FTE+SC co-culture did not further enhance ciliated differentiation (Figure 6C; FTE+SC+E2, purple line), despite the formation of even larger organoids (Figure 6A-B). Instead, lineage analysis revealed that E2 treatment led to reduced enrichment of multiple ciliated cell signatures (Ciliated_1, Ciliated_4, and Ciliated_3) and increased enrichment of Unclassified_1 and Unclassified_2 signatures (Figure 6C, dashed arrows, and Figure S6B). This reduction in ciliated differentiation was corroborated by co-IF staining, which showed reduced numbers of AcTUB^+^ cells in FTE+*PcY* SC+E2 organoids compared with FTE+*PcY* SC organoids (Figure 2C), and by qRT-PCR demonstrating reduced *Foxj1* expression in FTE+*PcY* SC+E2 organoids (compared to FTE+*PcY* SC organoids, Figure S2D). In contrast, addition of E2 (1nM) to FTE cells cultured without stromal cells (FTE+E2) did not elicit similar effects, but instead modestly increased several lineage signatures, including Unclassified_1, Secretory_1, Ciliated_4, and Ciliated_1 (Figure 6C, green line, and Figure S6C). Together, these data suggest that E2 acts primarily through FT stromal cells to increase the stemness of FTE cells, potentially by expanding primitive progenitor-like populations (e.g., Unclassified_1), intermediate progenitors (e.g., Unclassified_2), and/or by attenuating stromal cell-induced ciliated differentiation.

**Figure 6.**
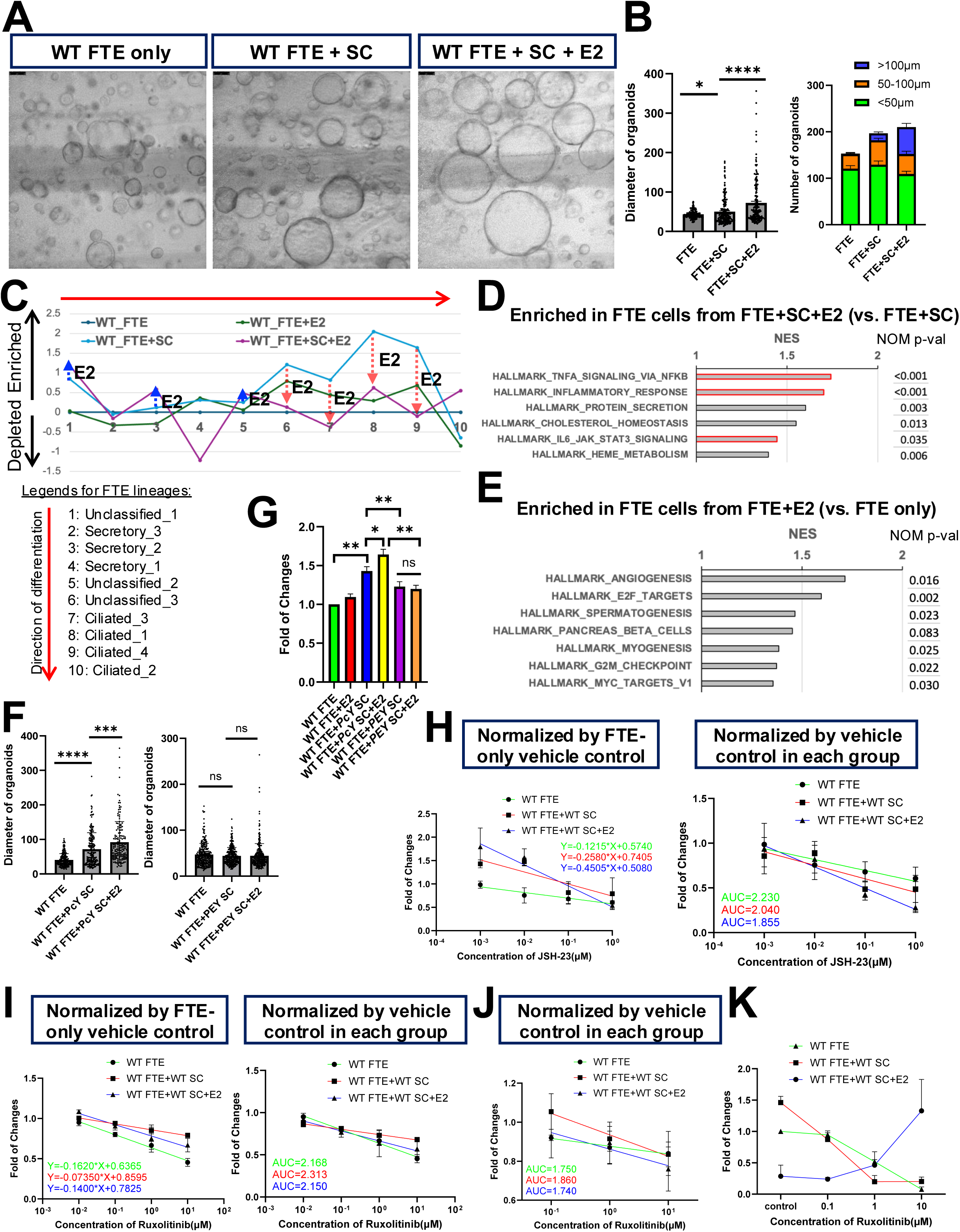
Estrogen promotes inflammation and growth of FTE cells via *Esr1^+^* stromal cells. (A) Representative pictures from organoid cultures for established WT FTE organoid cells growing in the BET medium alone (FTE only) or co-cultured with WT FT stromal cells (SC) with (FTE+SC+E2) or without (FTE+SC) Estradiol (E2) treatment (1nM). Scale bars=50μm. (B) Quantification of sizes and numbers (at different size ranges) of organoids formed from different cultures as in (A). *: p≤0.05, ****: p≤0.0001, Student’s t-test. Error bar represents ±SEM. (C) Relative enrichment or depletion (based on GSEA) of the indicated gene sets for FTE lineages (#1-10) based on human single-cell dataset in *Dinh et al* ^19^ in FTE organoid cells from the indicated cultures in relation to those from the WT FTE only culture (as baseline, =0). Arrows indicate E2-induced FTE lineage changes in FTE+SC+E2 cultures in relation to the FTE+SC cultures. (D-E) GSEA data showing top enriched gene sets in the Hallmark collection from the MSigDB in FTE cells from the FTE+SC+E2 cultures in relation to those from the FTE+SC cultures (D), or in FTE cells from the FTE+SC cultures in relation to those from the FTE only cultures (E). Gene sets related to inflammation/immune pathways are highlighted. (F) Quantification of sizes of organoids formed from established WT FTE organoid cells co-cultured with MACS-sorted FT stromal cells from *PcY* (lef) or *PEY* (right) mice with or without E2 treatment (1nM). Representative organoid pictures are shown in Figure S6D. ***: p≤0.001, ****: p≤0.0001, ns = not significant, Student’s t-test. Error bar represents ±SEM. (G) ATP assay data for organoids in Figures 6F and S6D. *: p≤0.05, **: p≤0.01, by Student’s t-test. Error bar represents ±SEM. (H) Linear regression curves of ATP assay for organoids from the indicated culture conditions treated with different concentrations of JSH-23 for 7 days. AUC=area under the curve. (I) Linear regression curves of ATP assay for organoids from the indicated culture conditions treated with different concentrations of Ruxolitinib for 7 days. (J) Linear regression curves of ATP assay for organoids from the indicated culture conditions treated with different concentrations of Ruxolitinib for 3 days. (K) qRT-PCR analysis of *Foxj1* expression in organoids from J treated with different concentrations of Ruxolitinib for 3 days.

To investigate the molecular basis of this indirect E2 effect, we further analyzed RNA-seq data from FTE organoid cells cultured under these conditions. Compared with FTE+SC cultures, E2 treatment in the presence of stromal cells (FTE+SC+E2) resulted in significant upregulation of multiple inflammation/immune-related gene sets in FTE cells (Figure 6D, highlighted). Importantly, E2 treatment of FTE cells in the absence of stromal cells did not induce similar changes (Figure 6E), indicating that E2-driven inflammatory signaling in FTE cells is mediated indirectly via stromal cells.

To directly test the requirement for stromal *Esr1* in this process, we co-cultured WT FTE organoid cells with FT stromal cells sorted from either *PcY* (WT) or *PEY* (*Esr1*-null) mice. As expected, the growth-promoting effect of the co-cultured stromal cells was substantially diminished when *PEY* stromal cells were used (Figures 6F-G and S6D). Importantly, the additional growth enhancement induced by E2 in the FTE+SC condition was abolished in the presence of *Esr1*-deficient stromal cells (Figures 6F-G and S6D), demonstrating that stromal *Esr1* is required for E2-mediated indirect effects on FTE growth.

To further examine the link between E2-induced inflammatory signaling and increased FTE stemness, we focused on the NFκB and JAK/STAT pathways, both of which have been implicated in promoting epithelial stemness ^38–40^. We treated organoid cultures with JSH-23 (an NFκB inhibitor) or Ruxolitinib (a JAK/STAT inhibitor)^41^ at increasing concentrations and assessed organoid growth using an ATP-based assay. Over a 7-day culture period, organoids under all conditions (FTE only, FTE+SC, and FTE+SC+E2) exhibited dose-dependent growth inhibition; however, organoids cultured under the FTE+SC+E2 condition showed the most pronounced reduction in growth in response to either inhibitor (Figure 6H-I, left plots, and Figure S6E). When each culture condition was analyzed as an independent “cell line” (i.e., for the purpose of testing their individual drug sensitivity) and growth was normalized to their corresponding vehicle-treated controls (=1), area under the dose-response curve (AUC) analysis revealed that FTE cells in the FTE+SC+E2 condition exhibited the greatest sensitivity to both NFκB and JAK/STAT inhibition (i.e., the lowest AUC values; Figure 6H-I, right plots). Consistent results were obtained using an alternative NFκB inhibitor, BMS (Figure S6F). These findings indicate that FTE cells exposed to E2 in the presence of stromal cells are more dependent on inflammatory signaling pathways to sustain their enhanced proliferation and as a result, they also exhibit a higher sensitivity to the inhibition of either NFκB or JAK/STAT pathway.

Finally, to assess the relationship between inflammatory signaling and ciliated differentiation, we measured *Foxj1* expression in FTE organoid cells under these different culture conditions in the presence or absence of pathway inhibitors. After 3 days of Ruxolitinib treatment, FTE organoids cultured under the FTE+SC+E2 condition already exhibited increased sensitivity relative to those cultured under the FTE+SC condition, as reflected by the lower AUC value (Figure 6J). Consistent with earlier observations, stromal co-culture alone increased *Foxj1* expression, whereas addition of E2 to the co-culture reduced *Foxj1* expression, indicating reduced ciliated differentiation under the FTE+SC+E2 condition compared to the FTE+SC condition (Figure 6K, control). Notably, Ruxolitinib treatment led to a dose-dependent upregulation of *Foxj1* expression specifically in the FTE+SC+E2 condition (Figure 6K), suggesting that JAK/STAT pathway activity contributes to the suppression of ciliated differentiation induced by E2-stimulated stromal signaling. In contrast, NFκB inhibition with JSH-23 did not produce a similar effect on *Foxj1* expression. Together, these data support a model in which estrogen exposure, acting through *Esr1^+^* FT stromal cells, indirectly activates inflammatory signaling pathways in FTE cells, thereby enhancing their stem-like properties. Increased activity of both NFκB and JAK/STAT pathways promotes FTE proliferation, while elevated JAK/STAT signaling specifically contributes to reduced ciliated differentiation.

### Stromal estrogen signaling promotes inflammation and growth of *Trp53/Brca1/Pten*-null FTE cells

To assess the potential impact of estrogen exposure on early cancer initiation from FTE cells, we examined how E2 treatment and FT stromal cells influence mutant FTE cells lacking the tumor suppressors *Trp53*, *Brca1*, and *Pten* (hereafter referred to as *PBP*-null FTE cells). *PBP*-null FTE cells have been shown to progress to HGSOCs following a relatively long latency ^13,42,43^. We found that compared with WT FTE cells, *PBP*-null FTE cells formed organoids that were more compact, solid, and dense in appearance (“PBP FTE” in Figure 7A compared to “WT FTE only” in Figure 6A). RNA-seq and lineage analysis using human FTE signatures defined by *Dinh et al*^19^ revealed that *PBP*-null FTE cells were largely blocked in ciliated differentiation, exhibiting increased enrichment of the Unclassified_3 signature, reduced ciliated cell signatures, and upregulation of cell cycle-associated gene sets (Figure 7B-C). Similar to WT FTE cells, treatment of *PBP*-null FTE organoids with varying concentrations of E2 alone did not result in significant changes in organoid size or morphology (Figure S7A). In contrast, co-culture of *PBP*-null FTE cells with WT FT stromal cells resulted in the formation of larger, more translucent organoids, a phenotype that was further enhanced by addition of E2 (1nM) (Figure 7A,D). Consistently, ATP-based assays demonstrated increased organoid growth in response to stromal co-culture, which was further augmented by E2 treatment (Figure 7E). IF staining confirmed that stromal co-culture increased the number of Ki67^+^ proliferative cells in *PBP*-null FTE organoids, and that E2 addition further enhanced this proliferative response (Figure S7B). RNA-seq analysis of sorted *PBP*-null FTE cells from these different culture conditions revealed that, relative to WT FTE cells (WT_FTE), all *PBP*-null conditions (±E2, ±stromal cells) shared similar global transcriptional changes driven by *PBP*-loss (Figure S7C). However, when *PBP*-null FTE cells cultured alone (PBP_FTE) were used as the baseline, E2 treatment alone exerted minimal effects (Figure 7F, orange line). In contrast, stromal co-culture expanded the Unclassified_1 subset and partially induced ciliated differentiation, as indicated by upregulation of the Ciliated_1, Ciliated_4, and Ciliated_2 signatures (Figure 7F, green line). However, addition of E2 (1nM) to the *PBP*-null FTE+SC co-culture reduced ciliated differentiation (i.e., downregulation of Ciliated_1 and Ciliated_4) and increased enrichment of the Secretory_1 signature (Figure 7, light blue line, and Figure S7D), suggesting estrogen exposure alters the lineage composition of *PBP*-null FTE cells indirectly through signaling via *Esr1*^+^ stromal cells, favoring a more secretory and less ciliated state.

**Figure 7.**
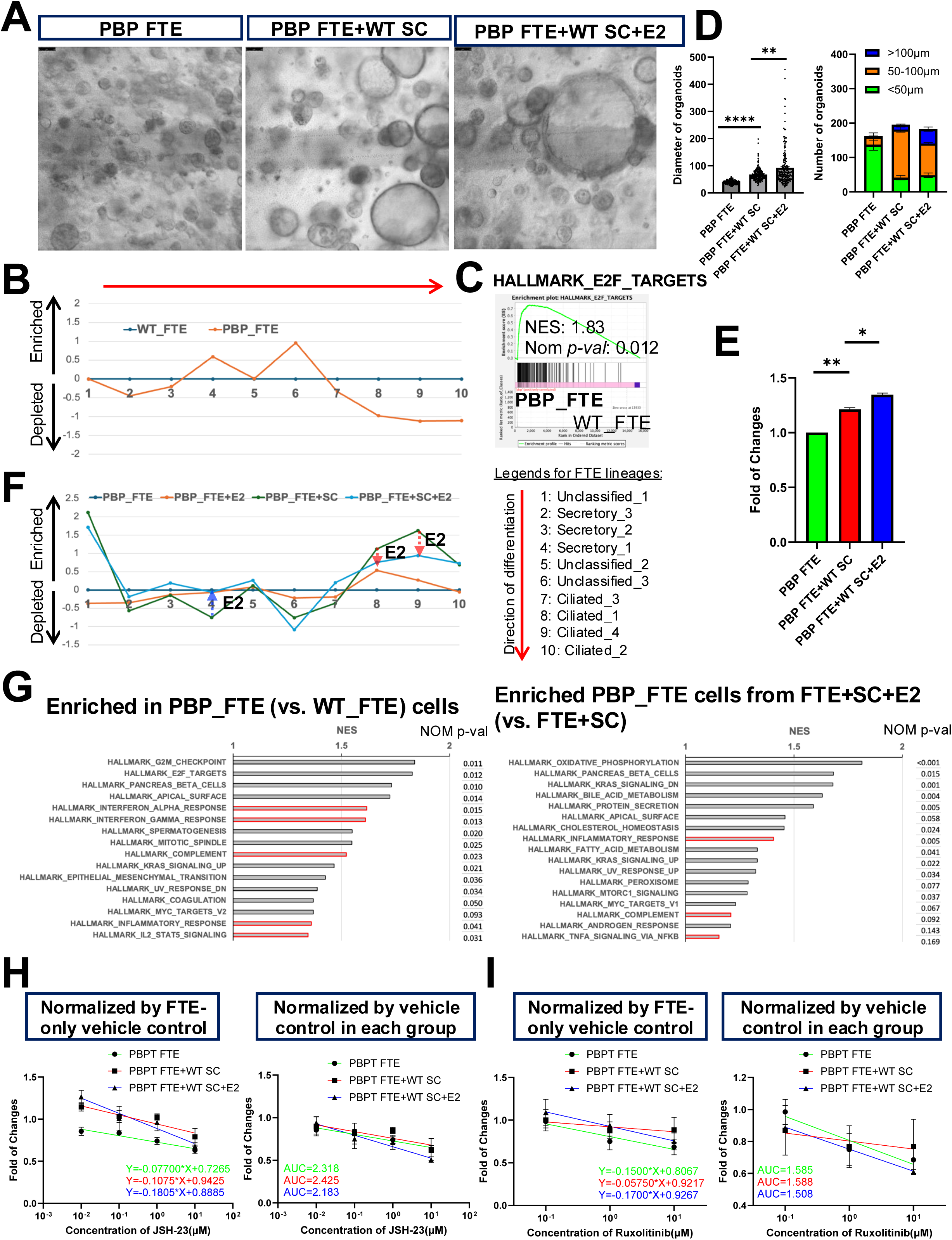
Estrogen promotes proliferation of *PBP*-null FTE cells via *Esr1^+^* stromal cells. (A) Representative pictures from organoid cultures for established *PBP*-null FTE organoid cells growing in the BET medium alone (FTE only) or co-cultured with WT FT stromal cells (SC) with (FTE+SC+E2) or without (FTE+SC) E2 treatment (1nM). Scale bars=50μm. (B) Relative enrichment or depletion (based on GSEA) of the indicated gene sets for FTE lineages (#1-10) based on human single-cell dataset in *Dinh et al* ^19^ in *PBP*-null FTE organoid cells (PBP_FTE) in relation to WT FTE organoid cells (WT_FTE) (as baseline, =0). (C) GSEA showing enrichment of the cell cycle-related Hallmark gene set in *PBP*-null FTE organoid cells (PBP_FTE) in relation to WT FTE organoid cells (WT_FTE). (D) Quantification of sizes and numbers (at different size ranges) of organoids in (A). **: p≤0.01, ****: p≤0.0001, Student’s t-test. Error bar represents ±SEM. (E) ATP assay of organoids in (A). *: p≤0.05, **: p≤0.01, Student’s t-test. Error bar represents ±SEM. (F) Relative enrichment or depletion (based on GSEA) of the indicated gene sets for FTE lineages (#1-10) based on human single-cell dataset in *Dinh et al* ^19^ in *PBP*-null FTE organoid cells from the indicated cultures in relation to those from the *PBP*-null FTE only cultures (as baseline, =0). Arrows indicate E2-induced FTE lineage changes in FTE+SC+E2 cultures in relation to the FTE+SC cultures. (G) GSEA data showing top enriched gene sets in the Hallmark collection from the MSigDB in *PBP*-null FTE cells in relation to WT FTE cells (left) and in *PBP*-null FTE cells from the FTE+SC+E2 culture in relation to those from the FTE+SC culture (right). Gene sets related to inflammation/immune pathways are highlighted. (H) Linear regression curves of ATP assay for organoids from the indicated culture conditions treated with different concentrations of JSH-23 for 7 days. AUC=area under the curve. (I) Linear regression curves of ATP assay for organoids from the indicated culture conditions treated with different concentrations of Ruxolitinib for 7 days.

We next examined inflammation/immune-related signatures in *PBP*-null FTE cells under these different conditions. Compared with WT FTE cells, *PBP*-null FTE organoids exhibited enrichment of multiple inflammation/immune-related gene sets (Figure 7G, highlighted), consistent with their increased proliferation and impaired differentiation. Inflammatory gene signatures were further upregulated in *PBP*-null FTE cells from FTE+SC+E2 cultures compared with FTE+SC cultures, whereas E2 treatment alone did not induce similar changes (Figures 7G and S7E), indicating that estrogen also enhances inflammatory signaling in *PBP*-null FTE cells indirectly through stromal cells. The magnitude of this induction was less pronounced than that observed in WT FTE cells (Figure 6D), likely reflecting the elevated basal inflammatory state of *PBP*-null FTE cells (Figure 7G).

To determine whether E2-induced inflammatory pathways contribute to the enhanced growth of *PBP*-null FTE cells, we treated their corresponding organoid cultures with inhibitors of NFκB (JSH-23) or JAK/STAT (Ruxolitinib). As observed in WT FTE cells, increasing concentrations of either inhibitor resulted in the most pronounced growth inhibition in *PBP*-null FTE organoids cultured under the FTE+SC+E2 condition (Figure 7H-I, left plots). AUC analysis confirmed that *PBP*-null FTE cells in the FTE+SC+E2 condition exhibited the greatest sensitivity to both inhibitors (i.e., lowest AUC values; Figure 7H-I, right plots). Together, these results indicate that estrogen exposure enhances proliferation and inflammatory signaling in *PBP*-null FTE cells primarily through *Esr1^+^* FT stromal cells, and that activation of NFκB and JAK/STAT pathways contributes to this stromal-mediated effect, potentially facilitating early stages of tumor initiation.

## Discussion

In this study, we uncover a previously unappreciated developmental role for estrogen/ER signaling in FT stromal cells in maintaining FT homeostasis. Although *Esr1* is expressed in both FTE cells and FT stromal cells, our findings indicate estrogen/ERα signaling serves distinct functions in these two compartments. In FTE cells, estrogen-epithelial ERα signaling has been shown to play a critical role during pregnancy by supporting fertilization and preimplantation embryo development through suppression of oviductal protease activity, as reported previously ^32^. Loss of *Esr1* in FTE cells does not appear to directly disrupt the homeostasis of the oviduct (FT) directly. Instead, our data reported here demonstrate that estrogen signaling through stromal ERα plays a dominant role in sustaining FT homeostasis, primarily by regulating FTE cell proliferation and ciliated differentiation (Figures 1-2). Consistent with this model, *in vivo* deletion of *Esr1* in FT stromal cells resulted in a marked reduction in FT size accompanied by reduced ciliated cell differentiation (Figure 1). Complementary *in vitro* co-culture experiments further showed that *Esr1*-WT stromal cells robustly promoted both proliferation and ciliated differentiation of FTE organoid cells, whereas *Esr1*-deficient stromal cells exhibited a substantially reduced capacity to support these processes (Figure 2). Together, these findings establish stromal ERα signaling as a key regulator of epithelial homeostasis in the FT.

Previous studies using human FT (hFT) organoids reported that estrogen treatment promotes FTE differentiation toward the ciliated lineage by downregulating *WNT7A*, a factor required to maintain FT stem cells ^44^. In our murine FTE-only organoid cultures, E2 treatment alone similarly elicited a modest increase in ciliated differentiation, as revealed by RNA-seq-based lineage analysis (Figure 6C, green line). However, the presence of FT stromal cells induced a far more pronounced differentiation of secretory cells toward the ciliated lineage (Figure 6C, light blue line), underscoring the dominant role of stromal-derived signals in this process. Notably, the concentration of E2 used in the hFT organoid study (100nM) was substantially higher than that employed here (1nM), which may contribute to the differences in the magnitude and context of estrogen responses observed between these systems.

At the molecular level, our data demonstrate that estrogen signaling through stromal ERα, but not epithelial ERα, regulates the expression of genes encoding inflammatory cytokines, growth factors, and ECM components (Figures 3 and S3). Using the FTE organoid system, we further showed that several of these stromal-derived inflammatory cytokines and growth factors can directly influence FTE cells by promoting their proliferation and/or differentiation toward the ciliated lineage (Figures 4-5). Intriguingly, although treatment of FTE cells with either E2 or FT stromal cells alone modestly or robustly induced ciliated differentiation, respectively, combined treatment with E2 and stromal cells resulted in reduced ciliated differentiation despite an overall increase in FTE organoid growth (Figure 6). We propose that this paradoxical effect reflects an indirect action of E2 mediated through stromal cells, leading to enhanced stem-like properties of FTE cells. Specifically, E2 signaling via *Esr1^+^* stromal cells augments the production of secreted factors, including inflammatory cytokines, which in turn activate inflammatory signaling pathways in FTE cells (e.g., JAK/STAT, NFκB). Activation of these pathways promotes FTE proliferation while simultaneously altering lineage composition, favoring more primitive or progenitor-like cellular states (Figure 6C).

Notably, nearly all cytokines tested in our study (in Figures 4-5) can activate the NFκB pathway ^45–48^, which likely contributes to cytokine-induced FTE differentiation. In contrast, a more restricted subset of stromal-derived factors (e.g., IL6, interferons, CCL2) can activate the JAK/STAT pathway ^46,49,50^. Activation of this pathway is likely a key driver of self-renewing proliferation in more primitive FTE stem/progenitor populations, including secretory and undifferentiated states. This model is supported by our observation that inhibition of the JAK/STAT pathway, but not the NFκB pathway, in the FTE+SC+E2 condition (but not in FTE+SC or FTE-only cultures) led to increased expression of the ciliated differentiation marker *Foxj1* (Figure 6K). This effect is likely attributable to a reduction in the stem/progenitor cell pool resulting from blockade of JAK/STAT pathway-dependent self-renewal proliferation. Consistent with this interpretation, the JAK/STAT pathway has been well established as a central regulator of stem cell proliferation and survival across multiple tissues ^51^.

Our observations linking stromal estrogen signaling to inflammatory pathways and FTE lineage composition are highly relevant to the dynamic cellular remodeling of the FT during the estrous cycle. At ovulation, corresponding to estrus, fimbrial cells are exposed to follicular fluid containing locally high concentrations of estrogen ^35,36^. Through signaling in ERα^+^ stromal cells, this estrogen surge may initiate a cascade of inflammatory responses analogous to those observed during wound repair. Activation of inflammatory pathways is expected to stimulate proliferation of FTE stem/progenitor cells, particularly within the secretory lineage, and to promote their differentiation in order to replenish epithelial cells damaged by the proinflammatory follicular fluid encountered during ovulation. Concomitantly, stromal production of inflammatory cytokines and growth factors is likely to drive ciliated differentiation, thereby increasing the abundance of mature ciliated cells required for efficient transport of the ovulated oocyte. In FT stromal cells, we identified *Pgr*, encoding the progesterone receptor, as a downstream target of estrogen signaling (Figures 3D and S3B). During the estrous cycle, estrogen levels peak during proestrus in mice, followed by a rise in progesterone levels during metestrus and diestrus ^34^. In contrast to estrogen-driven stromal signaling, progesterone signaling in stromal cells is thought to exert anti-inflammatory effects ^52^. Thus, activation of progesterone signaling may contribute to the resolution of estrogen-induced inflammatory responses and associated epithelial remodeling, thereby restoring FT homeostasis as the estrous cycle progresses and preparing the tissue for the subsequent cycle.

Our findings also have important implications for understanding aging-associated changes in the FT. FT aging has been linked to increased immune infiltration, which is thought to result from repeated ECM remodeling during successive estrous cycles and incomplete resolution of inflammation ^33^. Recent single-cell studies further indicate that inflammatory programs are markedly enhanced in FT stromal fibroblasts with aging, including upregulation of multiple inflammatory cytokines ^33^. Our results suggest that stromal-derived cytokines, exemplified by CCL2, play a functional role in maintaining epithelial homeostasis by supporting FTE proliferation and differentiation (Figure 5). The observation that aging-associated cytokine expression is estrogen dependent under homeostatic conditions, yet remains elevated in aged tissues despite declining estrogen levels, points to the emergence of estrogen-independent regulatory mechanisms. One potential explanation is the establishment of a persistent inflammatory state, or “inflammatory memory,” within FT stromal fibroblasts. Similar phenomena have been described in fibroblasts from chronic inflammatory diseases, where stable epigenetic remodeling sustains inflammatory gene expression independently of the initiating stimulus ^53,54^. Such persistent stromal inflammation during aging may alter epithelial-stromal interactions, disrupt FT homeostasis, and potentially create a detrimental microenvironment that predisposes the aging FT epithelium to pathological transformation.

Under physiological homeostasis, FT stromal cells promote differentiation of FTE cells toward the ciliated lineage, thereby functioning as a largely tumor-suppressive niche. How this stromal niche transitions from a tumor-suppressive to a tumor-promoting state, and how such a shift increases the risk of malignant transformation in FTE cells, has remained poorly understood. Our findings identifying stromal estrogen signaling as a regulator of FTE behavior through inflammatory pathways provide a mechanistic framework for this functional switch. During normal reproductive cycling, estrogen levels fluctuate, and subsequent activation of progesterone signaling is thought to attenuate inflammation in the FT. In contrast, our FTE+SC+E2 organoid system models a condition of sustained estrogen exposure acting through stromal cells. The increased stemness of FTE cells (i.e., increased proliferation, reduced ciliated differentiation) observed under this condition suggest that excessive or prolonged estrogen signaling may promote early stages of tumor initiation from FTE cells by converting the stromal niche from a differentiation-promoting, tumor-suppressive environment into one that favors stemness and proliferation. Specifically, our data support a model in which excessive estrogen exposure enhances self-renewal of FTE stem/progenitor cells by driving chronic stromal inflammation and activating stemness-associated inflammatory pathways, such as JAK/STAT, within the epithelium. In the context of female reproductive history, factors such as early menarche, late menopause, or nulliparity increase the number of lifetime menstrual cycles and cumulative estrogen exposure. These conditions may therefore contribute to the establishment of a persistent inflammatory state within the FT stroma, ultimately creating a more inflamed, tumor-promoting niche with advancing age.

The FT stromal niche may also transition from a tumor-suppressive to a tumor-promoting role through cooperation with genetic or epigenetic alterations in adjacent FTE cells. Our findings indicate that loss of key tumor suppressors (*Trp53*, *Brca1*, and *Pten*) in *PBP*-null FTE cells disrupts their differentiation capacity (Figure 7B), rendering them less responsive to stromal cues that normally promote maturation. Although stromal cells retain some ability to induce ciliated differentiation in these mutant FTE cells (Figure 7F), this effect is markedly attenuated relative to WT epithelium (Figure S7C). In contrast, stromal cells remain effective in stimulating proliferation of *PBP*-null FTE cells, an effect that is further amplified by estrogen exposure (Figures 7E and S7B). Moreover, sustained estrogen signaling (via stromal cells) counteracts even the modest differentiation-promoting influence of the stromal niche (Figure 7F), shifting the balance toward proliferation further. Together, these observations support a model in which incomplete resolution of stromal inflammation, potentially driven by repeated reproductive cycles or prolonged estrogen exposure, creates a toxic microenvironment that favors epithelial proliferation. In the presence of genetic or epigenetic lesions that impair epithelial differentiation, such a microenvironment may promote clonal expansion of aberrant FTE cells and thereby increase susceptibility to malignant transformation.

A tumor-supportive stromal niche termed high-risk mesenchymal stem cells (hrMSCs) was recently identified in human FTs ^55^. These epigenetically altered MSCs are enriched in *BRCA1/2* mutation carriers, increase with age, and are present in FTs prior to STIC formation. Functionally, hrMSCs promote DNA damage and enhance survival of FTE cells, and can drive their malignant transformation *in vivo* ^55^. hrMSCs express human MSC marker such as CD90 (*THY1*), CD73, and CD105, as well as WT1, a key transcription factor implicated in the tumor-promoting function of hrMCSs ^55^. Notably, *Esr1^+^* FT stromal cells in our study are *Pdgfra*(CD140a)^+^ cells, and CD140a is a recognized marker of mouse MSCs ^56^. Interrogation of our previously published FT single-cell RNA-seq dataset^25^ revealed that *Thy1* and *Wt1* are predominantly expressed within the *Pdgfra^+^Esr1^+^* stromal cell subpopulation. These observations raise the possibility that *Pdgfra^+^Esr1^+^* FT stromal cells share a common developmental origin with hrMSCs and may represent precursors to hrMSCs. Moreover, our findings suggest that estrogen signaling could influence the emergence or tumor-supportive functions of hrMSCs, thereby linking hormonal regulation of the stromal niche to early events in FT carcinogenesis.

## STAR ★ METHODS

Detailed methods are provided in the online version of this paper and include the following:

## Key resources table

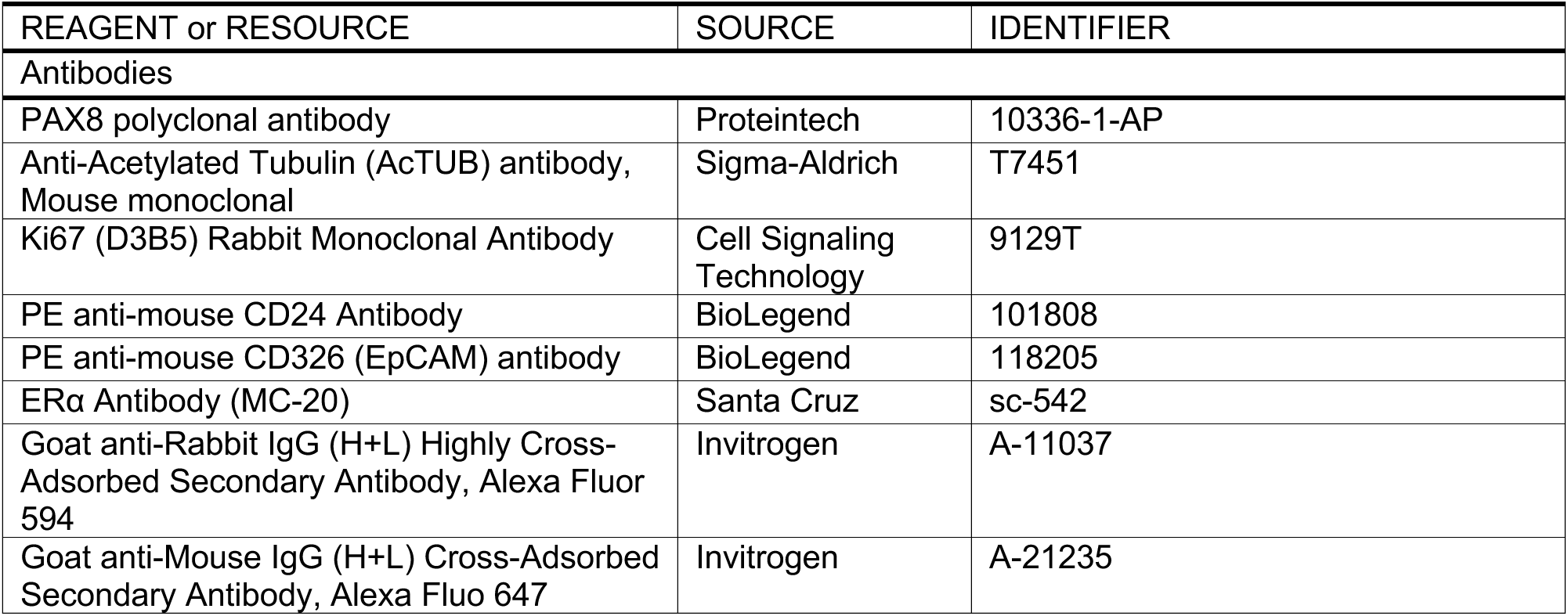
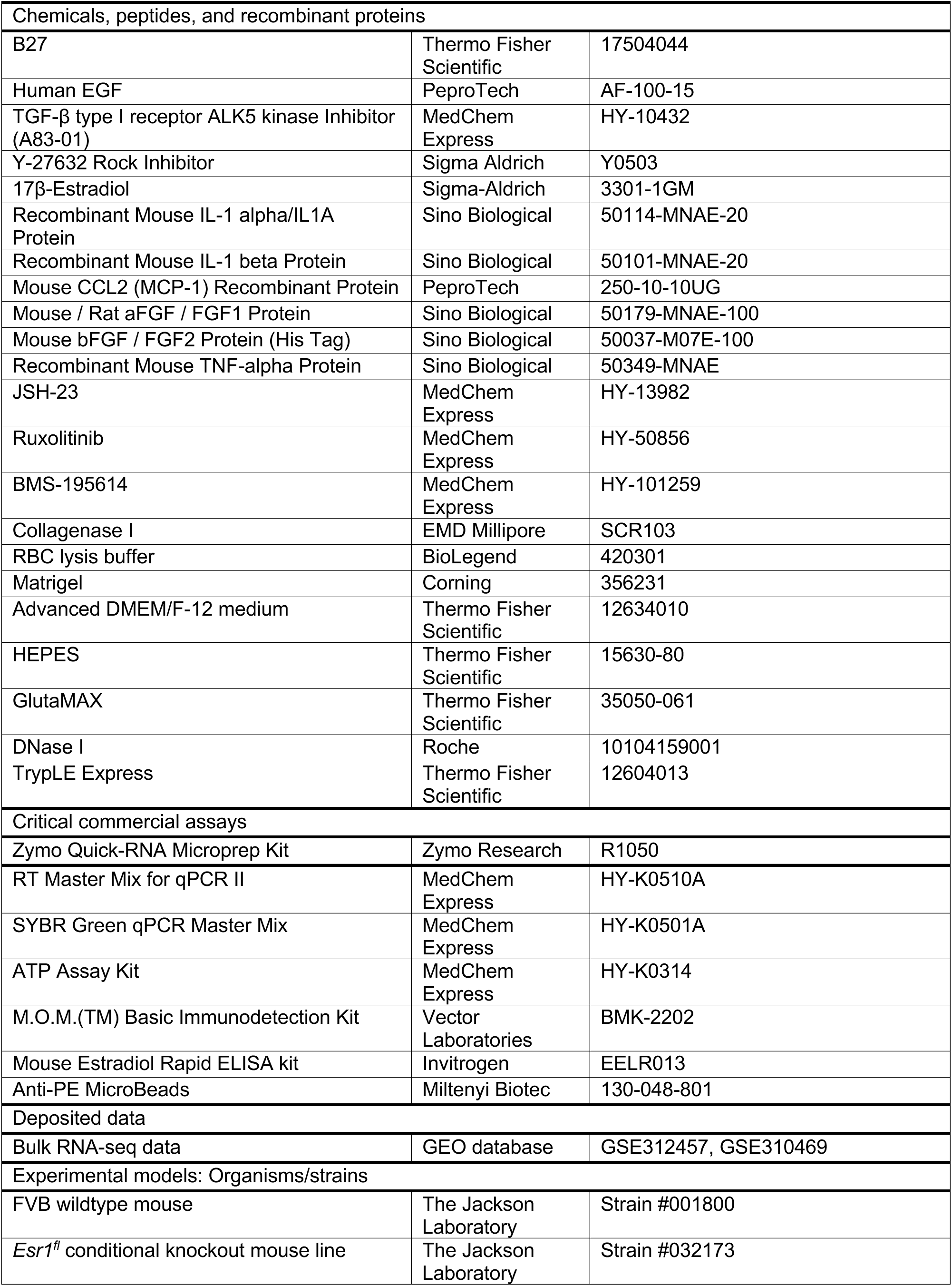
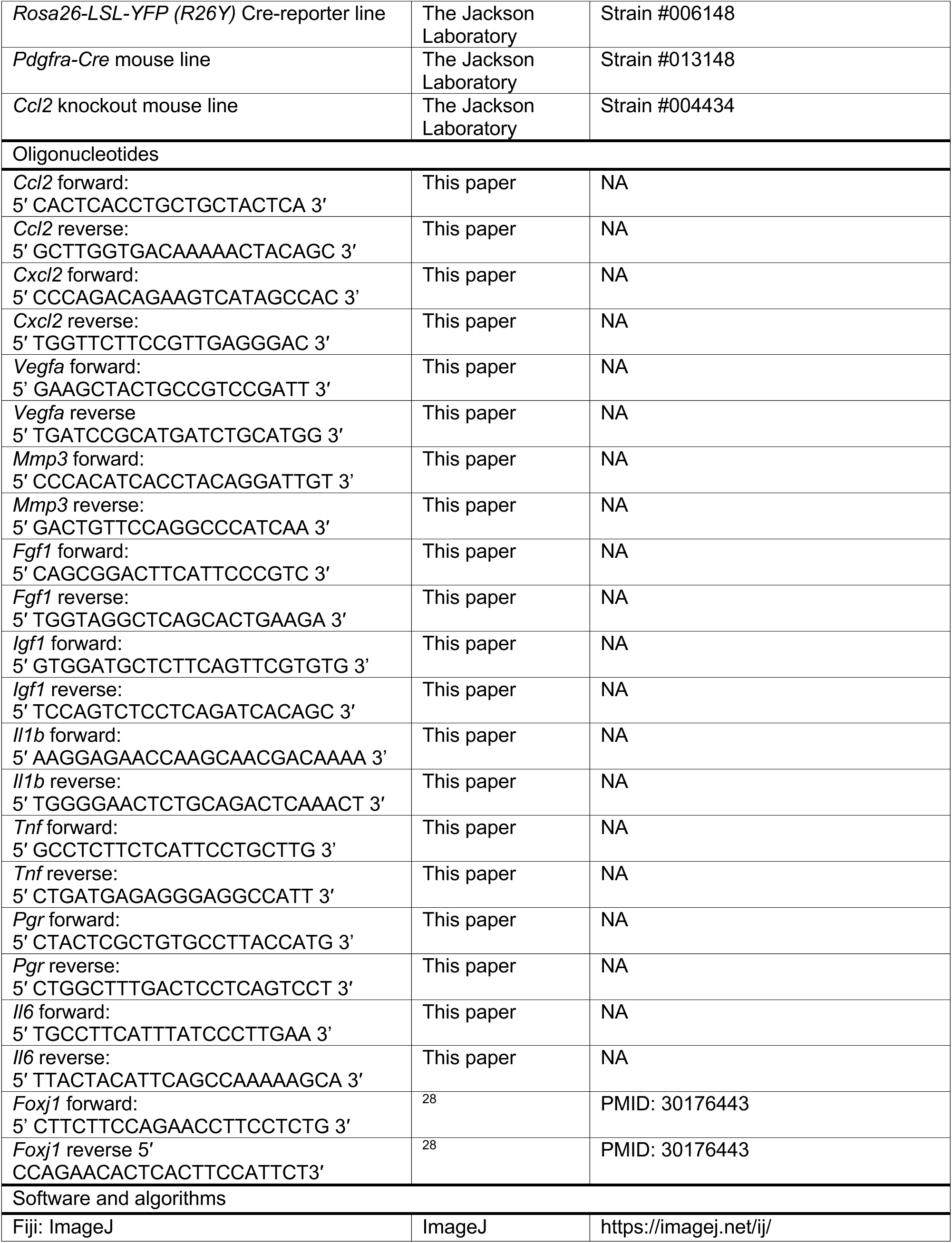
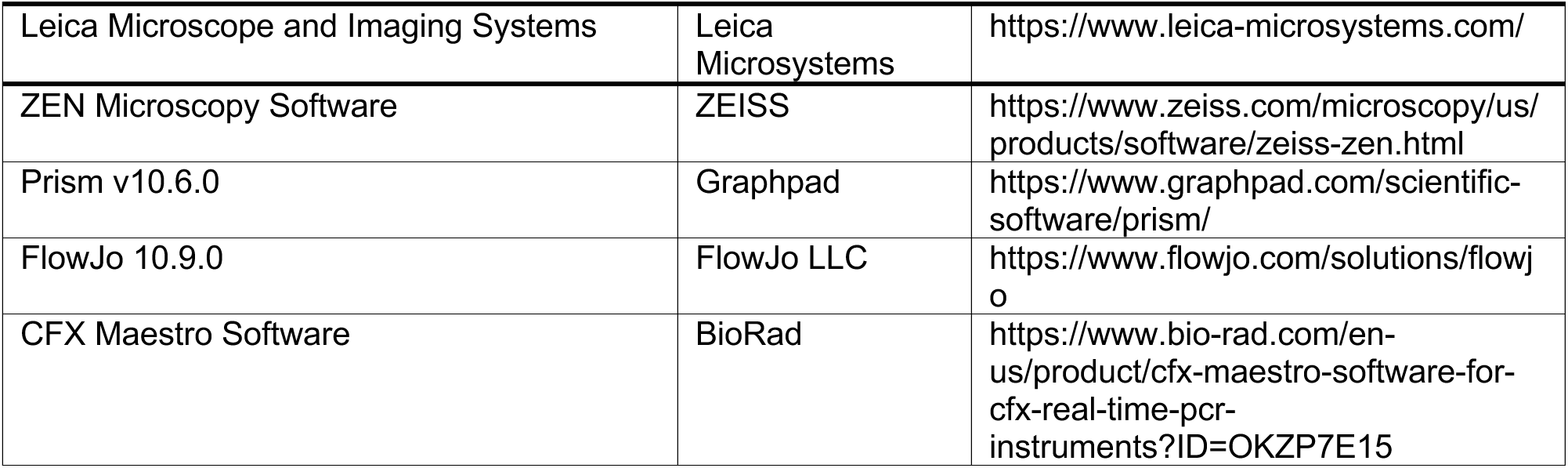

## Resource availability

### Lead Contact

Further information and requests for the resources and reagents should be directed to and will be fulfilled by the Lead Contact, Zhe Li (, or).

### Materials Availability

Further requests for resources and reagents should be directed to and will be fulfilled by the Lead Contact, Zhe Li (, or). This study did not generate new unique reagents.

### Data and Code Availability

The bulk RNA-seq dataset generated during this study has been deposited to Gene Expression Omnibus (GEO) repository (https://www.ncbi.nlm.nih.gov/geo/) with the dataset identifiers GSE312457 and GSE310469.

## Experimental model and subject details

### Mouse lines

All experimental procedures reported herein were reviewed and approved by our institutional animal care and use committee (IACUC) and performed in accordance with the relevant protocol (2023N000029). FVB stock mice [from The Jackson Laboratory (JAX) Strain #: 001800] were used as wild-type mice for FT organoid cultures. Mice entered the experiment between 8-12 weeks of age unless otherwise indicated. *Ccl2* knockout mice (JAX #: 004434), *Pdgfra-Cre* (JAX #: 013148) mice, *Esr1^fl^* (JAX #032173) conditional knockout mice, and the *Rosa26-LSL-YFP* (*R26Y*, JAX #: 006148) Cre-reporter mouse line were all purchased from JAX. *Pdgfra-Cre;Esr1^fl/fl^;R26Y* (*PEY*) experimental mice and *Pdgfra-Cre;R26Y* (*PcY*) control mice were generated by breeding *Pdgfra-Cre* mice with *Esr1^fl/fl^*and *R26Y* mice. *Ccl2^-/-^* (FVB) mice were obtained by backcrossing *Ccl2* knockout mice with FVB mice.

### Organoid culture and co-culture

Single-cell suspensions were embedded in 40 μl Matrigel (Corning #356231), resuspended, and allowed to solidify. The mixture was then overlaid with 400 μl pre-warmed BET or BETY organoid culture medium in one well of 12-well plate. BET medium refers to Advanced DMEM/F-12 medium (Thermo Fisher Scientific #12634010) with 10mM HEPES (Gibco^TM^ #15630-80), 1x GlutaMAX (Gibco^TM^ #35050-061), 100IU/ml of penicillin and 100μg/ml Streptomycin, 1x B27 (Thermo Fisher Scientific #17504044), 0.1μg/ml human EGF (PeproTech #AF-100-15), 0.5μM TGF-β type I receptor ALK5 kinase Inhibitor (A83-01) (MedChem Express # HY-10432). BETY medium refers to BET medium supplemented with 10μM Y-27632 Rock Inhibitor (Sigma Aldrich #Y0503), which is only necessary when organoids are first prepared (from fresh tissues) or thawed. Organoids were grown at 37 °C in a humidified atmosphere with 5% CO2. The medium was changed once every 3-5 days, and the organoids were passaged at a ratio of 1:4 once every 10-14 days. For passaging, the Matrigel containing the organoids was incubated in 1 ml TrypLE Express (Thermo Fisher Scientific #12604013) for 30 min, and then the single cell suspension was centrifuged at ∼350g for 5 min. Appropriate number of cells from the resulting pellet was resuspended in cold Matrigel and reseeded in the 12-well tissue culture plate.

The organoid/stromal cell co-culture was established by seeding 5,000 established FTE organoid cells (or freshly sorted FTE cells from FT tissues) and 5,000 MACS-sorted CD24^-^stromal cells using Anti-PE MicroBeads (Miltenyi Biotec #130-048-801) and PE anti-mouse CD24 Antibody (BioLegend #101808) from FT tissues of adult female mice with the indicated genotypes. The FTE cells either alone or mixed together with CD24^-^ stromal cells and Matrigel were seeded and cultured in the BET medium.

### Mouse stromal cells and FTE organoid sample preparation for bulk RNA-seq

To perform bulk RNA expression analysis of FT stromal cells, we isolated FT tissues from *Pdgfra-Cre;R26Y* and *Pdgfra-Cre;Esr1^fl/fl^;R26Y* adult female mice. To prepare FT cells as a single-cell suspension, FT tissues were dissected in dissection solution [PBS containing 2% fetal bovine serum (FBS) and 100IU/ml of penicillin and 100μg/ml Streptomycin]. After removing excessive connective and vascular tissues, the dissected FT tissues were minced into 1-2mm pieces with an iris scissor or forceps and then digested in 3.3mg/ml collagenase I (EMD Millipore #SCR103) and 1mg/ml DNase I (Roche # 10104159001) in DMEM/F12 medium for 1 hour at 37°C on a Nutating shaker. At the end of incubation, tissues were centrifuged at ∼350 g for 5 minutes, and the pellets were then incubated with 1x RBC lysis buffer (BioLegend #420301) for 3 minutes on ice. FT cells were then washed with 1x PBS and pelleted by centrifugation at ∼350 g for 5 minutes to achieve a single-cell suspension. YFP^+^ stromal cells were obtained by FACS sorting using a BD Aria II (561) sorter. Total RNA was isolated with Zymo Quick-RNA Microprep Kit (Zymo Research # R1050) according to the supplier’s protocol.

To isolate FTE cells from organoid culture or co-culture, the Matrigel containing the organoids was incubated in 1 ml TrypLE Express (Thermo Fisher Scientific #12604013) for 30 minutes at 37°C in a 5% CO2 incubator. and then the single-cell suspension was centrifuged at ∼350g for 5 min. The cells were stained with PE anti-mouse CD326 (EpCAM) antibody (BioLegend # 118205) for 20min and washed with 1x PBS. EpCAM^+^ FTE cells were obtained by FACS sorting using a BD Aria II (561) sorter. Total RNA was isolated with Zymo Quick-RNA Microprep Kit (Zymo Research # R1050) according to the supplier’s protocol.

### Bulk RNA-seq library preparation and sequencing

rRNA depletion was performed from 100ng of purified total RNA using QIAseq FastSelect rRNA HMR reagents according to manufacturer’s protocol. Libraries were prepared using Roche Kapa Biosystems RNA HyperPrep sample preparation reagents on a Beckman Coulter Biomek i7. Finished dsDNA libraries were quantified by Qubit fluorometer and Agilent TapeStation 4200. Uniquely dual indexed libraries were pooled in an equimolar ratio and shallowly sequenced on an Illumina MiSeq to further evaluate library quality and pool balance. The final pool was sequenced with paired-end 150bp reads on an Illumina NovaSeq X Plus at the Dana-Farber Cancer Institute Molecular Biology Core Facilities.

### Gene Set Enrichment Analysis (GSEA)

Visualization Pipeline for RNA-seq (Viper) analysis tool developed by Center for Functional Cancer Epigenetics (CFCE) at Dana-Farber Cancer Institute was used to generate standard outputs ^57^. For GSEA, Viper output data was normalized by using the DESeq2 module in GenePattern (https://www.genepattern.org/). GSEA was conducted by using the standalone GSEA program (https://www.gsea-msigdb.org/gsea/index.jsp) and the Molecular Signatures Database (MSigDB) (https://www.gsea-msigdb.org/gsea/msigdb), using the H: hallmark gene sets, C2: curated gene sets, and C5: ontology gene sets collections, or using gene sets for human FTE subpopulations extracted from FTE lineage signatures based on single-cell transcriptomics defined by *Dinh et al* ^19^. In this study, FTE lineage signatures were derived from genes differentially expressed across ten epithelial clusters defined by single-cell transcriptomics analysis and such genes were selected based on average log2 fold change (log2FC) > 0.5, adjusted p value < 0.01, with a requirement that at least 25% of cells within the cluster expressing the gene of interest ^19^. To calculate the relative enrichment of these FTE lineage signatures in FTE organoid cells sorted from FTE-only, FTE+SC, and FTE+SC+E2 cultures (i.e., Figures 6C, 7B, 7F, and S7C), a pairwise GSEA was performed for each FTE sample in relation to its corresponding baseline (i.e., WT_FTE in Figures 6C, 7B and S7C; PBP_FTE in Figure 7F); a reciprocal GSEA was also performed for each pair to account for any negative enrichment. The normalized enrichment scores from both comparisons for each pair were added as the relative enrichment value plotted on the Y axis.

### Immunofluorescence

The dissected FT tissues were fixed in 10% formalin and embedded in paraffin. Immunofluorescent labeling was carried out by following standard procedures, by incubating FT tissue section with primary antibody for PAX8 polyclonal antibody (Proteintech # 10336-1-AP, 1:200), Anti-Acetylated Tubulin (AcTUB) antibody, Mouse monoclonal (Sigma-Aldrich # T7451, 1:200), Ki-67 (D3B5) Rabbit Monoclonal Antibody (Cell Signaling Technology #9129T, 1:400), ERα Antibody (MC-20) (Santa Cruz #SC542, 1:100) diluted using M.O.M.(TM) Basic Immunodetection Kit (Vector Laboratories # BMK-2202); the section was then washed with 1x PBS, and incubated with the secondary antibody [Goat anti-Rabbit IgG (H+L) Highly Cross-Adsorbed Secondary Antibody, Alexa Fluor 594 (Invitrogen A-11037), Goat anti-Mouse IgG (H+L) Cross-Adsorbed Secondary Antibody, Alexa Fluo 647 (Invitrogen A-21235)] for 30 mins at room temperature.

### Drug Treatment and ATP Assay

Single-cell suspensions were embedded in 20 μl Matrigel (Corning #356231), resuspended, and allowed to solidify. The mixture was then overlaid with 250 μl pre-warmed BET or BETY organoid culture medium in one well of 24-well plate. 1nM 17β-Estradiol (Sigma-Aldrich #3301-1GM), JSH-23 (MedChem Express # HY-13982), BMS-195614 (MedChem Express # HY-101259) and Ruxolitinib (MedChem Express #HY-50856) were added to culture medium. Organoids were cultured for 3-7 days before processed for ATP release assay using ATP Assay Kit (MedChem Express #HY-K0314) according to the supplier’s protocol.

### RNA quantification and real-time PCR

Total RNA was isolated with Zymo Quick-RNA Microprep Kit (Zymo Research # R1050) according to the supplier’s protocol. Complete cDNA was synthesized from the isolated RNA by using RT Master Mix for qPCR II (MedChem Express # HY-K0510A). Real-time PCR was performed using the SYBR Green qPCR Master Mix (MedChem Express # HY-K0501A). PCR reaction was performed in triplicate. The amplification plots obtained from the qRT-PCR were analyzed with CFX Maestro Software (BioRAD); expression levels were quantified by applying the comparative C (threshold cycle) method and calculating ΔΔCt. Relative expression levels of the target genes were normalized to the expression of glyceraldehyde-3-phosphate dehydrogenase (*Gapdh* gene) in each individual sample.

### Quantification and statistical analysis

Leica Microscope and Imaging Systems and ZEN Microscopy Software were used for organoid and tissue section characterization and imaging. ImageJ was used for image processing and counting. All experiments were independently replicated at least three times. Statistical analysis was performed using Graphpad Prism v10.6.0, by Student’s t-test or by ANOVA followed by a Tukey’s analysis. Data was reported as Mean ± SEM unless otherwise indicated. Qualitative images presented are representative of the outcomes obtained in the replicate experiments.

## Acknowledgments

The authors would like to thank the Molecular Biology Core Facilities (MBCF) at Dana-Farber Cancer institute (DFCI), Boston, MA for assistance with processing and analyzing the RNA-seq data. The diagram in graphical abstract was created with BioRender.com. This research was supported by a U.S. Department of Defense Ovarian Cancer Research Program Pilot Award (HT94252310178) and Sundry Fund from Brigham and Women’s Hospital to ZL.

## Author contributions

ZL perceived the conceptual ideas, designed experiments, and supervised the project. XC, SH, DZ and GQ performed the experiments. XC and ZL analyzed the data and wrote the manuscript.

## Declaration of interests

The authors declare no competing interests.

## Supplemental Figure legends

**Figure S1. *Pdgfra-Cre*-induced *Esr1*-Loss leads to impaired fallopian tube (FT) homeostasis, related to Figure 1.**

(A) Dissection of ovary (OV), fallopian tube (FT) and uterus (UT) from 4-weeks old female mice with the genotype of control (*PcY*) or *Esr1* CKO (*PEY*). Scale bars=2.5 mm.

(B) Mouse estradiol (E2) concentration in the serum of estrous cycle stage-matched young adult female mice with the genotype of control (*PcY*) or *Esr1* CKO (*PEY*).

(C) Representative pictures of FT organoid cultures from *PcY* or *PEY* mice seeded with one FT per well. Scale bars=50μm.

(D) Quantification of sizes and numbers (at different size ranges) of organoids formed from the indicated organoid cultures as in (C). *: p≤0.05, Student’s t-test. Error bar represents ±SEM.

(E) Representative pictures of FT organoid cultures from *PcY* or *PEY* mice seeded with the same number of cells (1×10^4^, 5,000, or 1,000) per well. Scale bars=50μm.

(F) Quantification of sizes and numbers (at different size ranges) of organoids formed in the indicated organoid cultures as in (E). **: p≤0.01, Student’s t-test. Error bar represents ±SEM.

**Figure S2. Stromal *Esr1* is required for the regulation of FTE growth by stromal cells, related to Figure 2.**

(A) Schematic diagram showing how established FTE organoids are generated from Magnetic Activated Cell Sorting (MACS)-sorted CD24^+^ FTE cells and how the co-culture system is established from established FTE organoid cells and MACS-sorted FT stromal cells.

(B) GSEA data showing top enriched gene sets from the Biological Process (BP) subset of the C5: ontology collection from the Molecular Signatures Database (MSigDB) in FTE organoid cells sorted from the FTE cell/stromal cell co-cultures (FTE+SC) in relation to those from the FTE cell-only cultures (FTE). Gene sets related to ciliated cells are highlighted.

(C) Immunofluorescence (IF) staining of organoid sections for Ki67; arrows indicate organoid cells positive for Ki67. Scale bars= 50μm.

(D) Quantification of *Foxj1* expression levels in organoids from the indicated culture conditions (normalized to those from the FTE-only culture =1).

(E) Representative pictures of P0 organoids formed from the co-culture of freshly isolated WT FTE cells with FT stromal cells MACS-sorted from *PcY* or *PEY* mice (cultured for 7 or 14 days). Scale bars=50μm.

(F) Quantification of sizes and numbers (at different size ranges) of organoids formed in the indicated organoid cultures as in (E). *: p≤0.05, Student’s t-test. Error bar represents ±SEM.

**Figure S3. Stromal *Esr1* is linked with inflammatory pathways and secreted factors, related to Figure 3.**

(A) Volcano plot showing top differentially expressed genes in *Esr1*-null versus *Esr1*-wt YFP^+^ FT stromal cells from *PEY* versus *PcY* mice.

(B) Relative expression values of select genes in the indicated categories based on the RNA-seq data for *Esr1*-null (*Esr1*-ko) versus *Esr1*-wt YFP^+^ FT stromal cells from *PEY* versus *PcY* mice; *Gapdh* is included as a house-keeping gene to show no notable change in its expression upon *Esr1*-loss in FT stromal cells; error bars represent ± SEM.

(C) Relative expression values of select genes in the indicated categories based on the microarray data (based on GEO accession #: GSE37471) for *Esr1*-null (*Esr1*-ko) versus *Esr1*-wt oviduct (FT) tissues from pregnant female mice at day 1 or 2 of pregnancy; error bars represent ± SEM.

**Figure S4. Stromal *Esr1*-linked inflammatory cytokines and secreted growth factors regulate FTE growth, related to Figure 4.**

Co-IF staining of organoid sections for PAX8 and AcTUB. Organoids were derived from WT FTE cells growing in the BET medium supplemented with the indicated cytokines (as in Figure 4A). Arrows indicate organoid cells with apical AcTUB staining. Scale bars= 50μm.

**Figure S5. Aging-related stromal CCL2 regulates FTE growth, related to Figure 5.**

(A) Gene signatures at indicated estrous phases in FT (oviduct) fibroblasts from adult mice. P: proestrus; E: estrus; M: metestrus; D: diestrus. Data is from *Winkler et al* ^33^.

(B-C) Top upregulated genes (B) and Hallmark gene sets (C) in FT (oviduct) fibroblasts of 18-months old mice compared to those of young mice (in the diestrus phase, which is most similar to acyclicity). Data is from *Winkler et al* ^33^.

(D) IF staining of organoid sections for Ki67; arrows indicate organoid cells positive for Ki67. Scale bars= 50μm.

**Figure S6. Estrogen promotes inflammation and growth of FTE cells via *Esr1^+^* stromal cells, related to Figures 6.**

(A) Representative pictures from organoid cultures for established WT FTE organoid cells growing in the BET medium treated with different concentrations of Estradiol (E2). Scale bars=50μm.

(B) GSEA data showing depleted (left) or enriched (right) gene sets for FTE lineages based on human single-cell dataset in *Dinh et al* ^19^ in FTE organoid cells from the FTE+SC+E2 cultures in relation to those from the FTE+SC cultures.

(C) GSEA data showing enriched gene sets for FTE lineages based on human single-cell dataset in *Dinh et al* ^19^ in FTE organoid cells from the FTE+E2 cultures in relation to those from the FTE only cultures.

(D) Representative pictures of organoids formed from established WT FTE organoid cells co-cultured with MACS-sorted FT stromal cells from *PcY* (*Esr1*-wt) or *PEY* (*Esr1*-null) mice with or without E2 treatment (1nM). Scale bars=50μm.

(E) Representative pictures of organoids formed from established WT FTE organoid cells only, or co-cultured with MACS-sorted WT stromal cells with or without E2 treatment (1nM), treated with different concentrations of the indicated inhibitors. Scale bars=50μm.

(F) Linear regression curves of ATP assay for organoids from the indicated culture conditions treated with different concentrations of IKK inhibitor BMS-195614 for 7 days. AUC=area under the curve.

**Figure S7. Estrogen promotes proliferation of *PBP*-null FTE cells via *Esr1^+^* stromal cells, related to Figure 7.**

(A) Representative pictures from organoid cultures for established *PBP*-null FTE organoid cells growing in the BET medium treated with different concentrations of Estradiol (E2). Scale bars=50μm.

(B) IF staining of organoid sections for Ki67; arrows indicate organoid cells positive for Ki67. Scale bars= 50μm.

(C) Relative enrichment or depletion (based on GSEA) of the indicated gene sets for FTE lineages (#1-10) based on human single-cell dataset in *Dinh et al* ^19^ in *PBP*-null FTE organoid cells from the indicated cultures in relation to those from WT FTE only cultures (WT_FTE as baseline, =0).

(D) GSEA showing enriched gene sets for FTE lineages based on human single-cell dataset in *Dinh et al* ^19^ in *PBP*-null FTE organoid cells from the FTE+SC+E2 cultures in relation to those from the FTE+SC cultures (top) or in *PBP*-null FTE organoid cells from the FTE+SC cultures in relation to those from the FTE+SC+E2 cultures (bottom).

(E) GSEA data showing top enriched gene sets from the Hallmark collection from the MSigDB in *PBP*-null FTE cells from the FTE+E2 cultures in relation to those from the FTE only cultures.

## Notes

### Competing Interest Statement

The authors have declared no competing interest.

