## Supplementary figures and images for "Stromal estrogen signaling regulates fallopian tube homeostasis and cancer initiation via inflammatory pathways"

### Supplemental Figures S1-S7

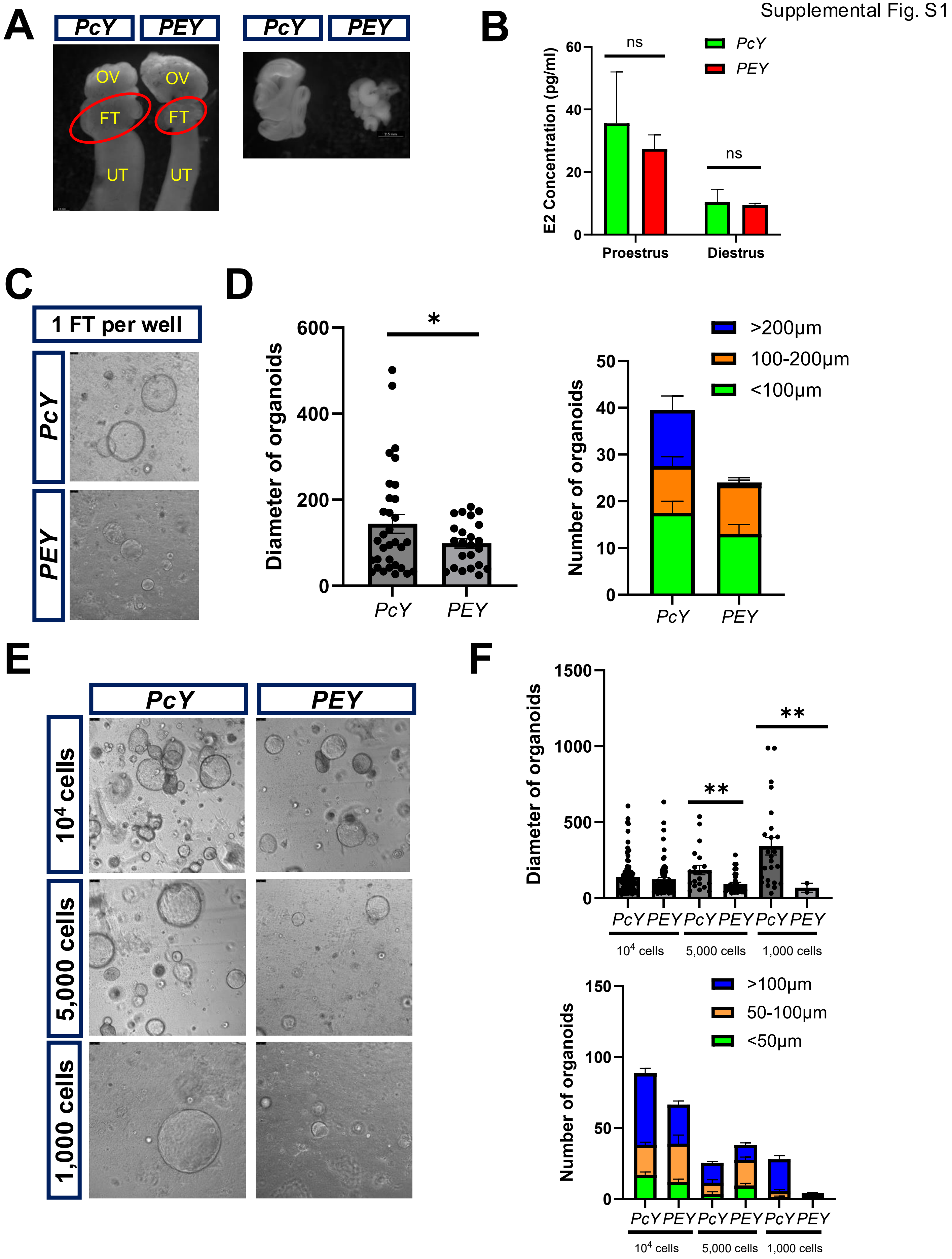

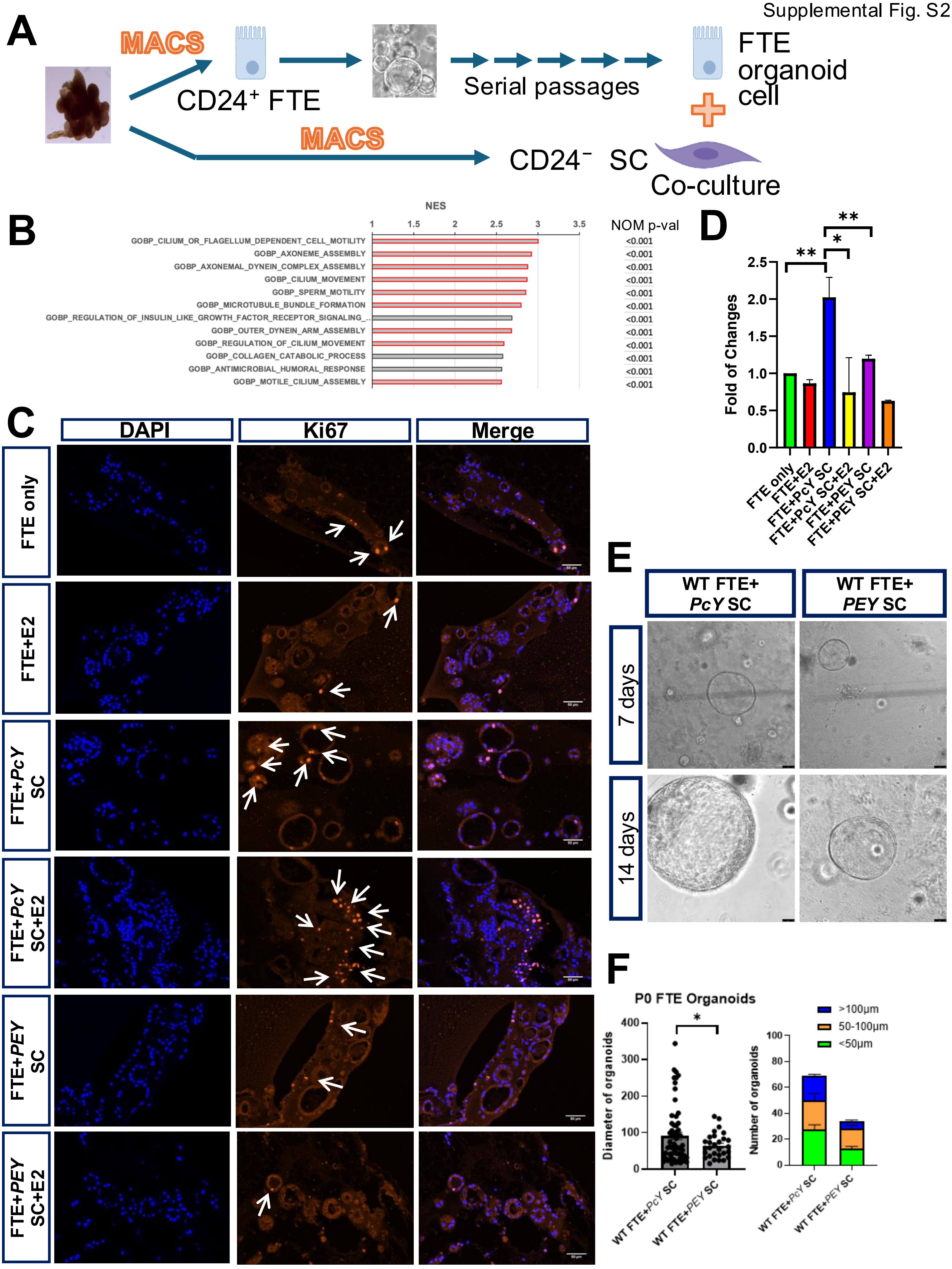

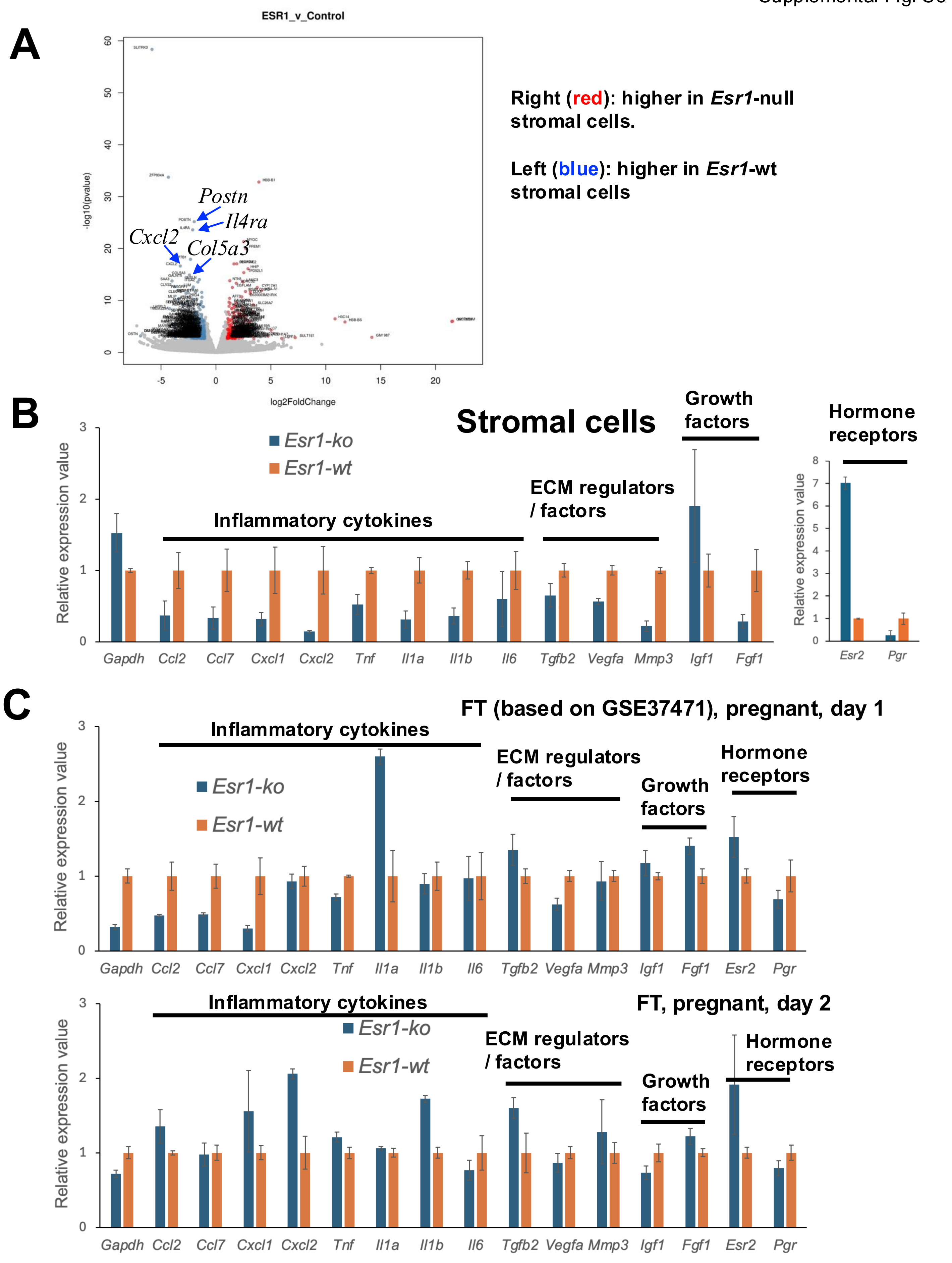

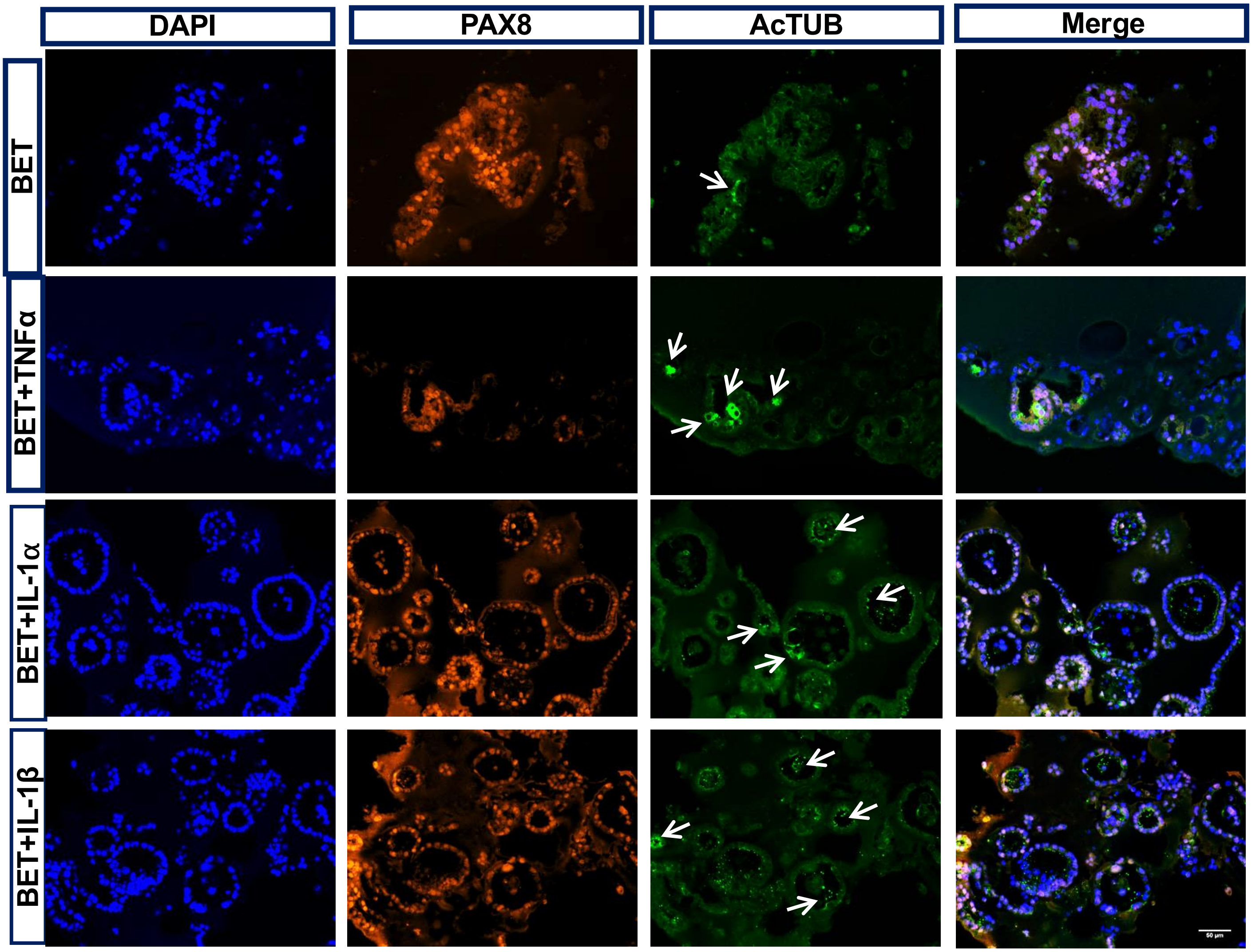

Supplemental Fig. S5

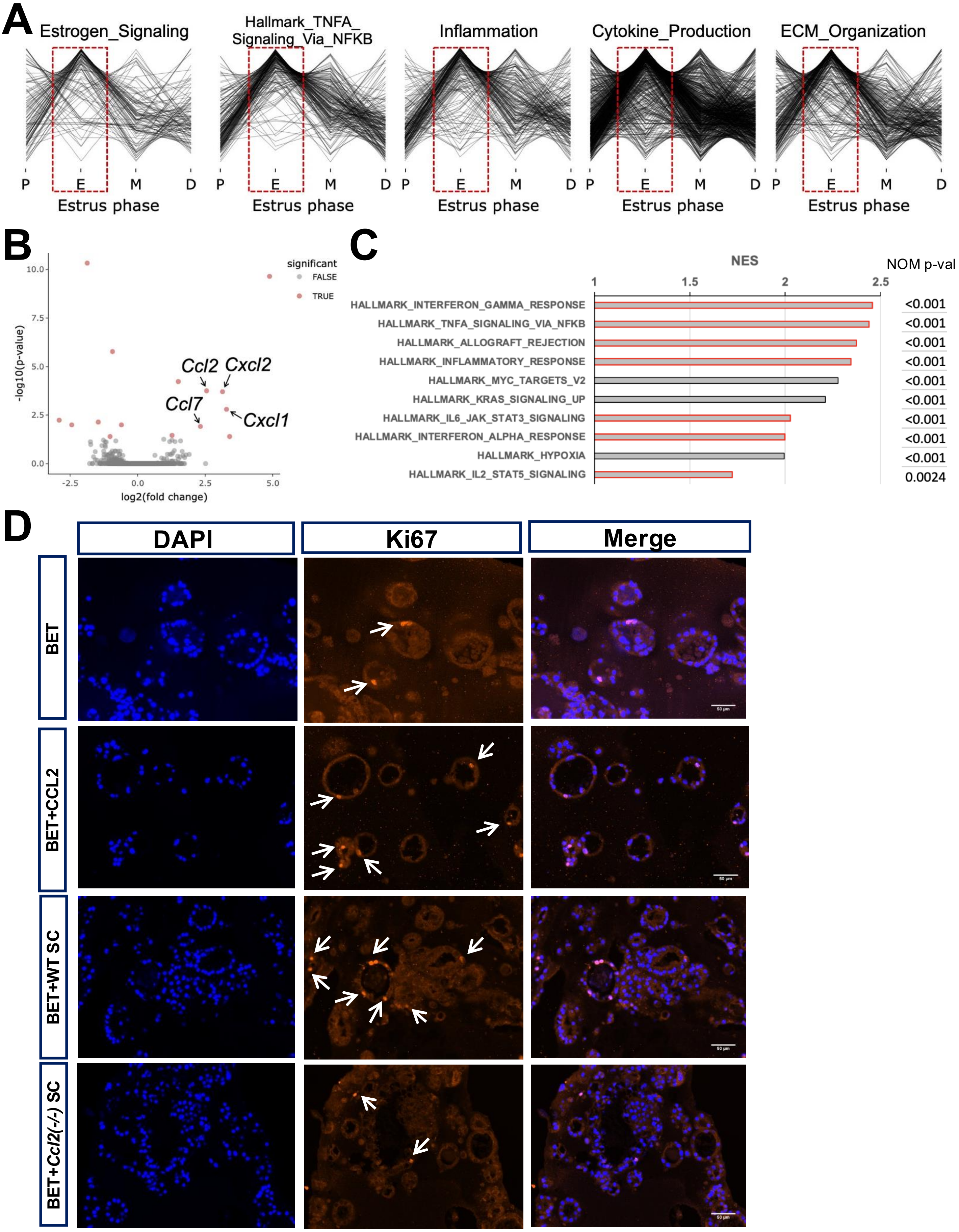

Supplemental Fig. S6

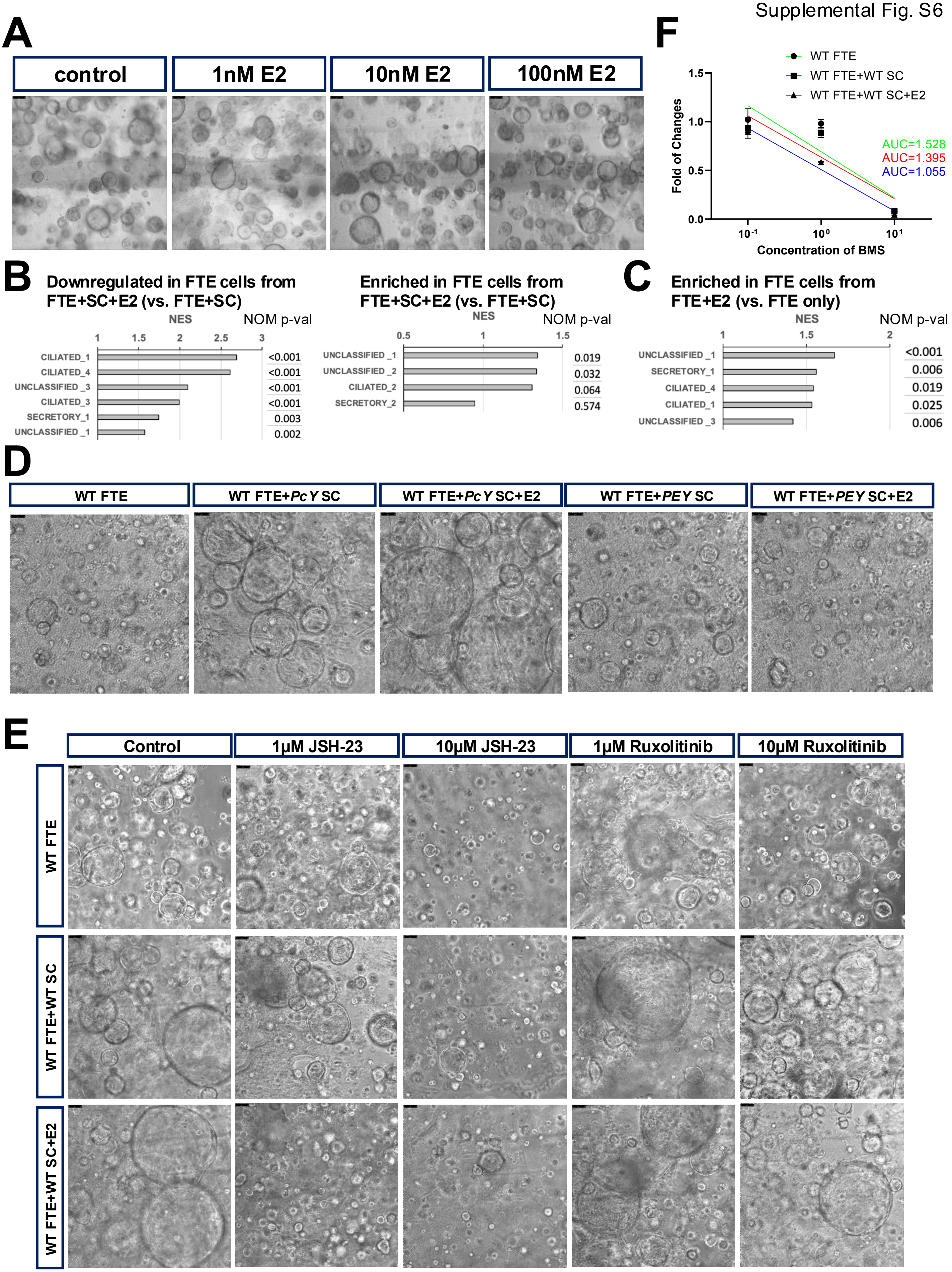

Supplemental Fig. S7

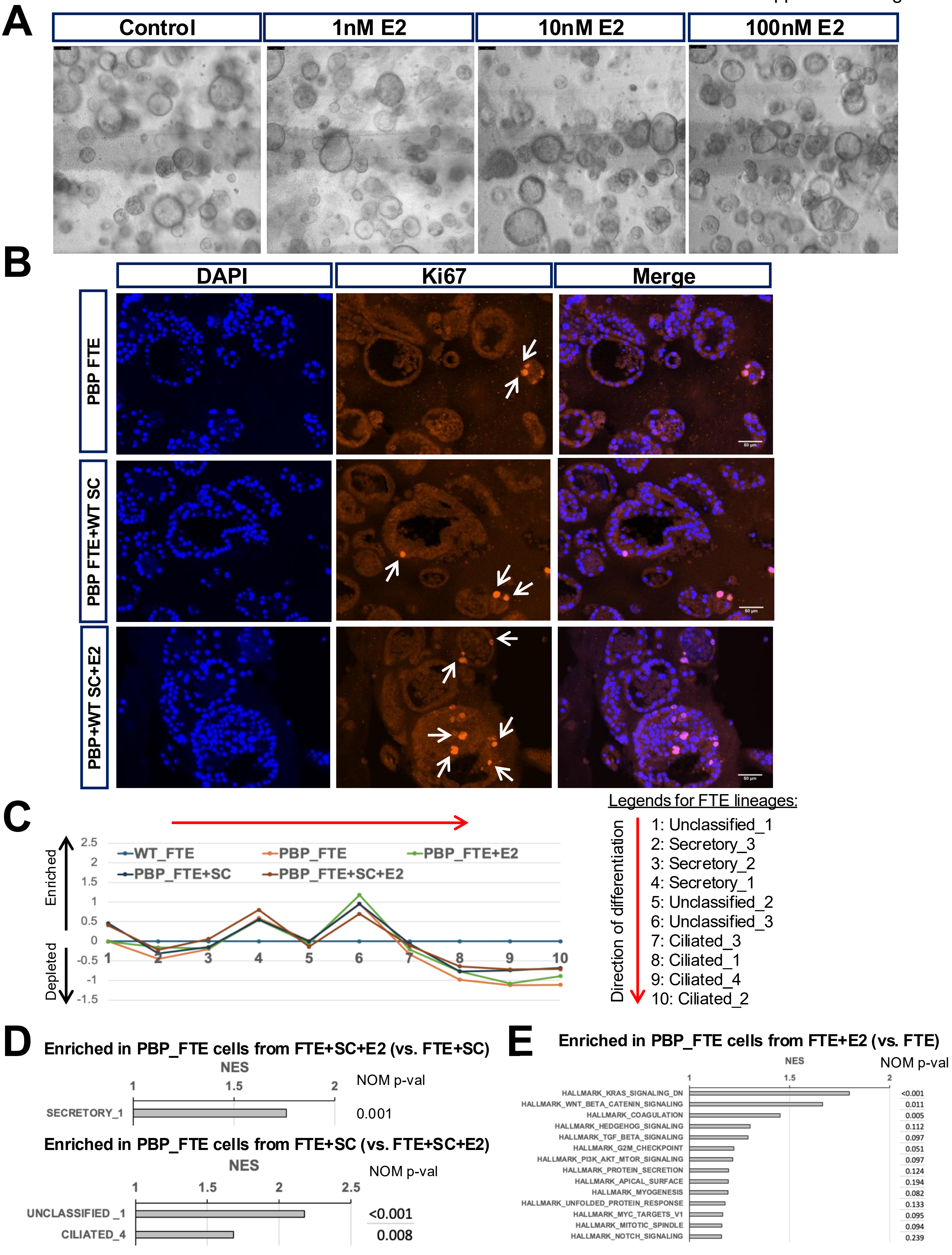
